# Millimeter-scale magnetic implants paired with a fully integrated wearable device for wireless biophysical and biochemical sensing

**DOI:** 10.1101/2023.11.23.568392

**Authors:** Ji Wan, Zhongyi Nie, Jie Xu, Zixuan Zhang, Shenglian Yao, Zehua Xiang, Xiang Lin, Yuxing Lu, Chen Xu, Pengcheng Zhao, Yiran Wang, Jingyan Zhang, Yaozheng Wang, Shaotong Zhang, Jinzhuo Wang, Weitao Man, Min Zhang, Mengdi Han

**Author notes:** These authors contributed equally: Ji Wan, Zhongyi Nie.

## Abstract

Implantable sensors can directly interface with various organs for precise evaluation of health status. However, extracting signals from such sensors must rely on transcutaneous wires, integrated circuit chips, or cumbersome readout equipment, which increases the risks of infection, reduces the biocompatibility, or limits the portability. Here, we develop a set of millimeter-scale, chip-less and battery-less magnetic implants that can measure biophysical and biochemical signals wirelessly. In particular, the implants form two-way communications with a fully integrated wearable device, where the wearable device can induce a large-amplitude damped vibration of the magnetic implants and capture their subsequent motions in a wireless manner. Such damped vibrations reflect not only the biophysical conditions surrounding the implants movements, but also the concentration of a specific biochemical depending on the surface modification. Experiments in rat models demonstrate the capabilities in measuring cerebrospinal fluid (CSF) viscosity, intracranial pressure (ICP), and CSF glucose levels. This miniaturized system opens possibility for continuous, wireless monitoring of a wide range of biophysical and biochemical conditions within the living organism.

## Introduction

Implantable sensors can measure a diverse array of information within the body (1–6), including electrophysiology (7–9), biomechanics (10–13), and concentrations of neurotransmitters and other biomarkers (14–16), to support the prevention and treatment of various diseases. These sensors exhibit changes in voltage, current, resistance, or capacitance to reflect the biophysical and biochemical conditions (17–20), but must rely on transcutaneous wires for data acquisition or signal transmission. In many cases that require long-term, continuous monitoring (e.g. epilepsy, diabetes, and heart failure) (21–24), implantable sensors should (1) transmit the signals outside the body in a wireless fashion, to avoid infection and inflammation caused by wires (25,26), and (2) feed the signals into data acquisition systems with miniaturized dimensions, to support measurement in settings outside of hospital or laboratory without bulky instruments (27).

However, current solutions to meet the above requirements rely on commercial chips, such as Bluetooth or near-field communication (NFC) system-on-chips (SoCs), that poses challenges in minimally invasive insertion, biocompatibility and power supply (28–30). Although implantable sensors based on inductor–capacitor (LC) resonant circuit and ultrasonic backscatter represent alternative solutions to reduce the dimensions of implantable sensors or to eliminate batteries, they demand bulky external acquisition systems (e.g. network analyzer, ultrasound probe, reading coil, lock-in electronics) that are not suitable for wearable applications (31–34) (text S1).

Here, we present a set of chip-less, battery-less magnetic implants with overall dimensions in millimeter scale. The implants can wirelessly communicate with a fully integrated centimeter-scale wearable device through magnetic field, to enable multimodal sensing of biophysical and biochemical signals. Key components of an implant involve a micro-magnet to generate alternate magnetic field during vibration, a soft, elastomeric membrane to improve the vibration amplitude, and surface coatings to selectively absorb targeted biochemicals. The chip-less and battery-less nature of the implant promotes its biocompatibility. In vivo experiments in rat models prove that the miniaturized system (i.e. the millimeter-scale implants and the centimeter-scale wearable device) can measure CSF viscosity, ICP and CSF glucose concentration continuously and wirelessly, with additional capability in real-time display on mobile terminals. This system opens avenues to long-term, continuous monitoring of a variety of signals related to health status, with potential to expand to wireless sensing of many other biochemicals and biomolecules through surface modifications.

## Results

### Working principle of the miniaturized system

As shown in Fig. 1A, we developed a miniaturized system that pairs a chip-less (35), battery-less magnetic implant in millimeter scale with a fully integrated wearable device (36) mounted at the skin surface. The wearable device can produce a pulsed magnetic field to wirelessly excite large-amplitude vibration of the magnetic implant. The subsequent damped vibration of the magnetic implant generates a dynamic magnetic field that can be captured by magnetic sensors integrated in the wearable device. These analog signals are then digitized and transmitted to a mobile terminal via Bluetooth for real-time display and further analysis (fig. S1). Specifically, the wearable part consists of a copper coil (wire diameter: 100 μm, turns: 1000) that converts a pulse current into a magnetic field, a tunneling magnetoresistance (TMR) sensor that measures the magnetic field generated by the magnetic implant, four Hall sensors for localization of the magnetic implant, a Bluetooth SoC that communicates with a mobile terminal wirelessly, a rechargeable lithium-ion battery for onboard power supply (capacity: 1200 mAh, voltage: 7.4 V), and a soft encapsulation layer in silicone (Fig. 1B). The magnetic implant includes a micro-magnet (diameter: 1.5 mm, height: 0.8 mm) coated with a layer of parylene-C (thickness: 10 μm), an elastic membrane in polydimethylsiloxane (PDMS, diameter: 6 mm, thickness: ∼40 μm), and a PDMS cavity substrate (diameter: 6 mm, height: 2 mm, cavity diameter: 5.5 mm, cavity height: 1 mm). The micro-magnet locates at the center of the elastic membrane, and the membrane bonds to the cavity substrate to allow for large-amplitude vibration under coil excitation (Fig. 1C and figs. S2 to S4).

**Fig. 1.**
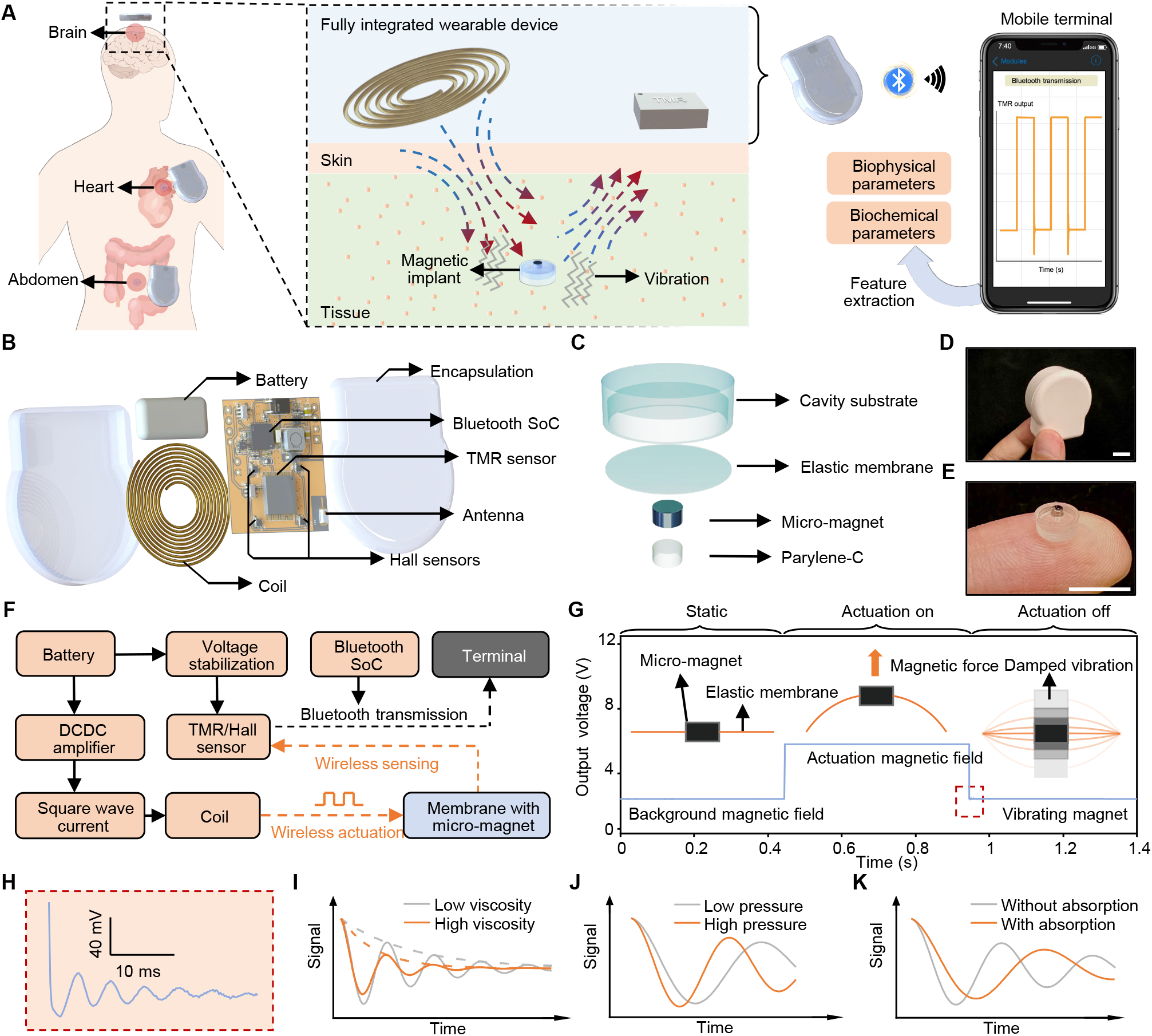
Design and working principle of the millimeter-scale magnetic implants and fully integrated wearable device. (**A**) Schematic illustrations of the two-way wireless communication between the millimeter-scale magnetic implants and the fully integrated wearable device, and the data acquisition and processing in mobile terminal. (**B** and **C**) Exploded schematic illustrations of the wearable device (B) and magnetic implants (C). (**D** and **E**) Optical images of the wearable device (D) and magnetic implants (E). Scale bars, 1 cm. (**F**) Block diagram of the circuit of the fully integrated wearable device for wireless actuation and sensing. (**G**) Schematic illustration of the motions of the micro-magnet, and corresponding vibration signals measured from the wearable device. (**H**) Magnified view of the vibration waveform depicted within the dashed red box in (G). (**I** to **K**) Schematic illustrations of the vibration waveforms under different viscosities (I), pressures (J), and concentrations of a specific chemical (K).

Figure 1 (D and E) outlines optical images of the centimeter-scale wearable device and the millimeter-scale magnetic implant. In the wearable device, a voltage stabilization module ensures a steady 7.4 V voltage for the TMR sensor, the Hall sensors, and the Bluetooth SoC. A direct current to direct current (DCDC) transformer boosts the battery output to 15 V for the coil, in order to provide sufficient magnetic field for exciting the magnetic implant. The Bluetooth SoC and an optocoupler switch modulate the 15 V voltage into a square wave with a frequency of 4 Hz. The coil converts this alternate voltage into a magnetic field with a peak intensity of 4.09 mT at a distance of 5 mm to excite the magnetic implant periodically (Fig. 1F, figs. S5 and S6 and movie S1).

During operation, a programmed alternating magnetic field generated from the wearable device periodically drags and releases the micro-magnet of the implant (Fig. 1G and figs. S7 and S8). The elastic and low-modulus membrane of the implant enable large-amplitude and non-linear damped vibration after release (Fig. 1H). The initial vibration amplitude reaches 0.4 mm under a pulse voltage of 15 V (distance between the wearable device and magnetic implant: 5 mm), two orders of magnitude larger those of conventional micro-electro-mechanical systems (MEMS) (37,38) (fig. S9). Such large vibration amplitudes create obvious variations of magnetic field at distances up to 9 mm, thereby facilitating wireless measurements in unshielded environments (fig. S10). As shown in Fig. 1H, a TMR sensor at a distance of 5 mm captures the variations of magnetic field wirelessly with a high signal-to-noise ratio (SNR, 28 dB of the first cycle), and produces a distinctive damped vibration signal upon the release of the magnetic implant without being affected by the presence of intermediate medium materials (fig. S11).

The large-amplitude, damped vibration signal comprises a rich set of information related to the physical and chemical environments surrounding the magnetic implant. First, viscosity holds immense value in the analysis of blood lipids and body fluids (39,40). For a magnetic implant with either a sealed or an open cavity (figs. S2 and S12), a surrounding environment with larger viscosity accelerates the attenuation of its vibrations (Fig. 1I). Second, pressure, such as ICP, intraocular pressure, blood pressure, and abdominal pressure, represent important parameters in the realm of implantable sensing. For a magnetic implant with a sealed cavity, an elevation in its surrounding pressure deforms the elastic membrane. This buckled membrane exhibits a stiffening effect, and leads to an increase in its vibration frequency (41,42) as depicted in Fig. 1J. Third, surface modifications on the parylene-C encapsulation or the elastic membrane enable selective binding of the target analyte through protein, nucleic acid or cells. As the concentration of a specific biomarker increases, more absorption occurs at the surface, leading to an increment of the mass of the membrane. This heavier membrane decreases the vibration frequency, as illustrated in Fig. 1K (43,44).

### Wireless, multiplexed sensing of physical and chemical conditions

To verify the illustrations in Fig. 1 (I to K), we perform finite element analysis (FEA) and experimental measurements of different magnetic implants designed for sensing of viscosity, pressure, and glucose concentration. Figure 2 (A and B) illustrates the vibrations of the magnetic implant in liquid environments with different viscosities. As viscosity increases, the damping force exerted on the micro-magnet becomes stronger, resulting in a faster decay rate. In the FEA model, altering the viscosity from 3.5 cP to 4.5 cP accelerates the attenuation of the vibration signal, with a fitted damping factor increasing from 56.68 to 81.47 (Fig. 2A, fig. S13A and movie S2). Experimental setup exploits mixtures of glycerol-water solutions with varying proportions to create liquid environments with different viscosities. Similar to the FEA, a mixture of 49% glycerol and 51% water (corresponding to a viscosity of 4.787 cP) enhances the damping of the magnetic implant (fitted damping factor: 100.53), compared with a mixture of 42% glycerol and 58% water (corresponding to a viscosity of 3.466 cP, fitted damping factor: 93.86, Fig. 2B and fig. S13B).

**Fig. 2.**
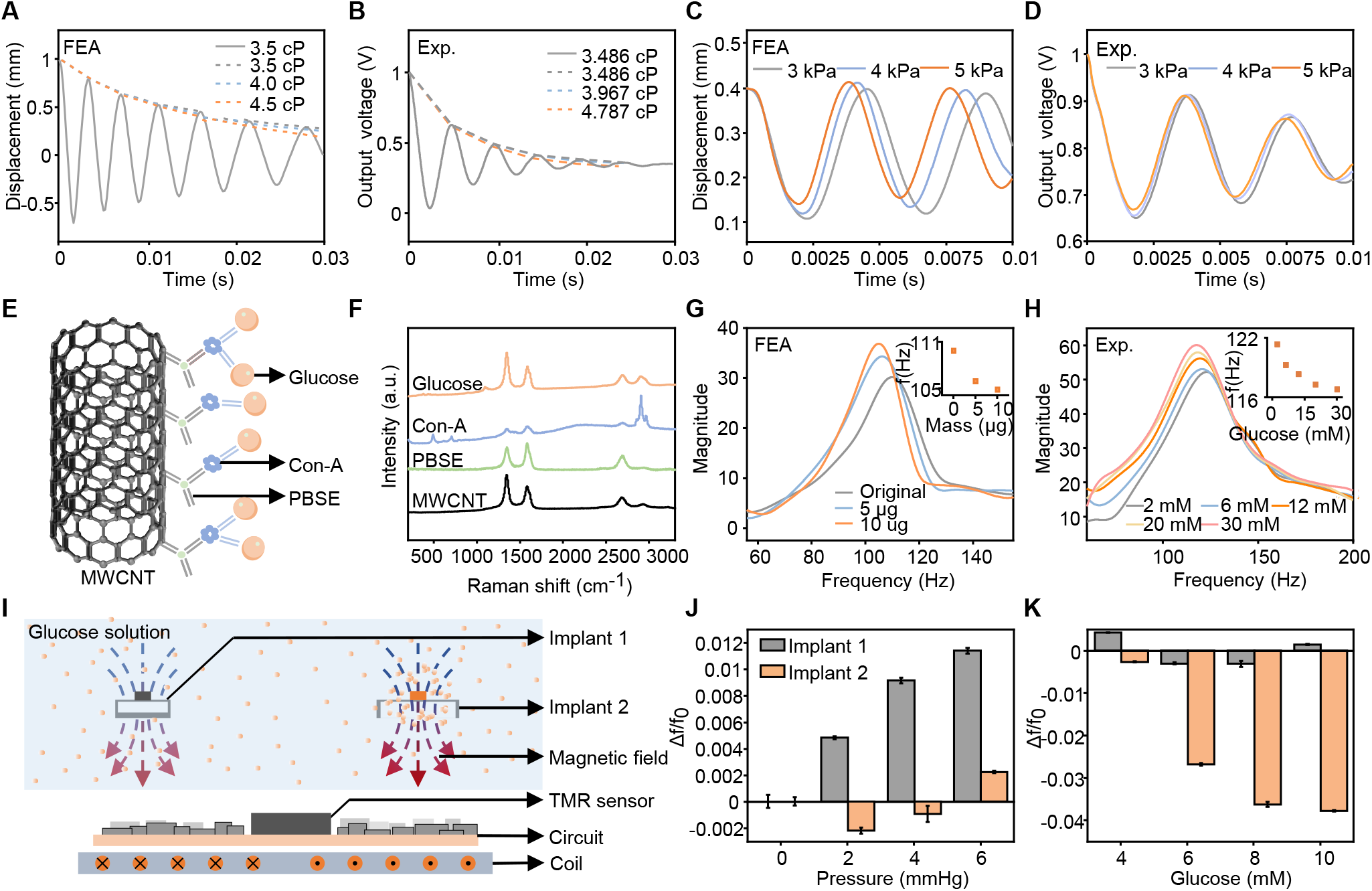
Responses of the magnetic implants under different physical and chemical conditions. (**A** and **B**) FEA predictions (A) and experimental measurements (B) of the vibrations of the magnetic implants (sealed cavity) under different viscosities. (**C** and **D**) FEA predictions (C) and experimental measurements (D) of the vibrations of the magnetic implant (sealed cavity) under different pressures. (**E**) Schematic illustration of the MWCNTs modified with 1-pyrenebutanoic succinimidyl ester-concanavalin A (PBSE-Con-A) for selective binding of glucose molecules. (**F**) Raman spectroscopy analysis of the Con-A modified MWCNTs. (**G** and **H**) FEA predictions (G) and measured vibrations (H) of the magnetic implant (open cavity) with different adhesion masses and different glucose concentrations (Fast Fourier transform, FFT of the vibration). The insets show the relationship between frequency and adhesion masses or glucose concentration. (**I**) Schematic illustration of two magnetic implants used for multiplexed sensing of pressure and glucose concentration. (**J** and **K**) Δf/f_0_ of the two magnetic implants for multiplexed sensing of pressure (J) and glucose concentration (K). Error bars correspond to the calculated standard deviation from five measurements.

Figure. 2 (C and D) shows the responses of a magnetic implant with a sealed cavity (internal pressure: 1 atm) under different external pressures. FEA results indicate that a pressure of 2 kPa deforms the membrane (thickness: 40 μm, diameter: 5.5 mm) downwards, with a displacement of 0.36 mm (Fig. 2C and fig. S14). The membrane under various pressures exhibits different effective stiffnesses and deformations. Therefore, pulling the micro-magnet upward by an external magnetic field (intensity: 4.09 mT at a distance of 5 mm) and releasing it by erasing the magnetic field result in a damped vibration of the micro-magnet along with the elastic membrane (movie S3). In experiments, we simulate the pressure of body fluids using liquid environment, and adjust the pressure through the height of the liquid level (fig. S15). The testing results reveal that a pressure increase from 3 kPa to 5 kPa raises the vibration frequency of the first cycle from 151.63 Hz to 160.51 Hz (Fig. 2D).

To achieve wireless sensing of glucose level, we employ chemical modifications on the surface of the micro-magnet to enable specific adsorption of glucose. The modification exploits a vertical dense array of multiwalled carbon nanotubes (MWCNTs) (45) to increase the overall surface area, thereby boosting the total amount of glucose absorbed on the micro-magnet (fig. S16). Concanavalin A (Con-A) modification on the MWCNTs enables specific binding of glucose (46) (Fig. 2E). The Raman spectra of the MWCNTs material revealed distinct peaks near 1345 cm-1 (D peak), 1573 cm-1 (G peak), and 2679 cm-1 (2D peak). Comparing the results to the unmodified MWCNTs, after PBSE modification, a peak emerged around 1623 cm-1, which can be attributed to the resonance of the pyrene moiety (47). This peak provides evidence of PBSE attachment to the MWCNTs. Furthermore, the presence of three new peaks near 2905.2 cm-1 after Con-A incubation indicates the successful attachment of Con-A (48) (Fig. 2F). Here, the magnetic implant adopts an open-cavity design to decouple the influence of pressure (fig. S17). Based on this magnetic implant, higher concentration of glucose leads to larger mass of the micro-magnet and lower vibration frequency. FEA results in Fig. 2G and movie S4 reveal a decrease in vibration frequency as the mass increases. Experiments in Fig. 2H and fig. S18 show a similar trend. As the glucose concentration varies from 2 mM to 30 mM, the vibration frequency decreases from 118.9 Hz to 116 Hz. The surface modification of Con-A endows a high selectivity to glucose over sodium ions, potassium ions, and lactic acid (fig. S19).

The above working principles underpin the capabilities of the magnetic implant in wireless sensing of viscosity, pressure, and glucose concentration. The elastic feature of the membrane enhances the amplitude of vibration, thereby creating a larger variation of the magnetic field to facilitate measurement using fully integrated, miniaturized wearable devices at unshielded environments. However, the large amplitude renders a non-linear feature of the vibration, and complicates the analytical analysis. Here, we utilize the multi-layer perceptron (MLP) regression algorithm in deep learning to ensure accurate calibration of the wireless sensors (fig. S20A). The feature set involves the time-domain waveform of the non-linear vibration and the frequency-domain waveform obtained through FFT (fig. S20B).

Using one cycle of damped vibration as the feature set, the sampling rate can achieve 15 Hz (i.e. actuating the micro-magnet 15 times per second, fig. S21). In order to mitigate the influence of noise, especially for the cases that use smaller magnets or require wireless acquisition at further distances, we exploit 16 consecutive cycles of damped vibrations as the test set (sampling rate to ∼1 Hz). The R2 of the model in this case reaches to over 95% (fig. S22A, 95.5% for viscosity, 99.89% for pressure, 98.1% for glucose concentration). As shown in fig. S22 (B to D), when providing the model with a substantial volume of test data (over 1000 cycles), the MLP regression models can predict the viscosity, pressure and glucose concentration with R2 surpassing 98%. The results indicate a trade-off between sampling rate and measurement accuracy.

Furthermore, vibration frequencies of the magnetic implants can be designed to cover a broad range (fig. S23 and text S2). As shown in fig. S24, an elastic circular membrane with a larger diameter leads to lower vibration frequencies. Optimization of the dimensions of the magnetic implants can tune their vibrations to separated frequency bands. As shown in fig. S25, a membrane (open cavity) with a thickness of 40 μm and a diameter of 8 mm exhibits a vibration frequency of 73 Hz, while another membrane (sealed cavity) with different dimensions (thickness: 40 μm, diameter: 6 mm) vibrates at a frequency of 100 Hz. The difference in frequency band creates possibilities for multiplexed wireless sensing using multiple magnetic implants. (49) As shown in Fig. 2I, experimental demonstration of multiplexed sensing incorporates two magnetic implants, including implant 1 (sealed-cavity, membrane thickness: 40 μm, membrane diameter: 6 mm, without surface modification) for the measurement of pressure, and implant 2 (open-cavity, membrane thickness: 40 μm, membrane diameter: 8 mm, with surface modification) designed for sensing of glucose concentration. This configuration enables simultaneous detection of liquid pressure and glucose concentration. The activation coil and the TMR sensor from the same wearable device can excite the vibrations of the two magnetic implants and record the variations of the magnetic field. The fractional change in frequency (Δf/f0) reveals that the implant 1 mainly responses to the pressure, whereas implant 2 shows a more obvious dependence to glucose concentration (Fig. 2, J and K). These results demonstrate the capability of the magnetic implants in multiplexed wireless sensing.

### In vivo biophysical and biochemical sensing in rat models

The chip-less and battery-less nature, as well as the biocompatible encapsulation (i.e. parylene-C), ensure biosafety of the magnetic implants. As shown in Fig. 3 (A and B), proliferation experiment (co-cultured L929 rat fibroblast cells) on the magnetic implant shows similar results compared to the control group, indicating its minimal impact on cell proliferation. The biocompatibility makes the magnetic implants suitable for in vivo applications. Chronic implantation of the magnetic implants in rat brain (two weeks: n=4, 3 months: n=2, control group: n=1) are necessary to evaluate their long-term stability and biocompatibility. By utilizing rabbit anti-Iba1 (1:500, Abcam) to label microglia and mouse anti-chondroitin sulfate (1:400, Sigma) to detect production of chondroitin sulfate proteoglycan, immunofluorescence test on the rat brain (two weeks after implantation) confirms that the magnetic implants have no adverse effects on the brain tissue (Fig. 3C and figs. S26 and S27). Three months after implantation, hematoxylin-eosin staining (H&E staining) on multiple organs, including brain, heart, lung, liver, spleen and kidney, demonstrates that the implants do not trigger inflammation in the organs (Fig. 3D). The results indicate that the elements present in the magnetic implants do not disperse or infiltrate into other organs over an extended duration of implantation. The cell viability, immunofluorescence test and H&E staining together validate the biosafety of the magnetic implants, thereby proving their suitability for biomedical applications.

**Fig. 3.**
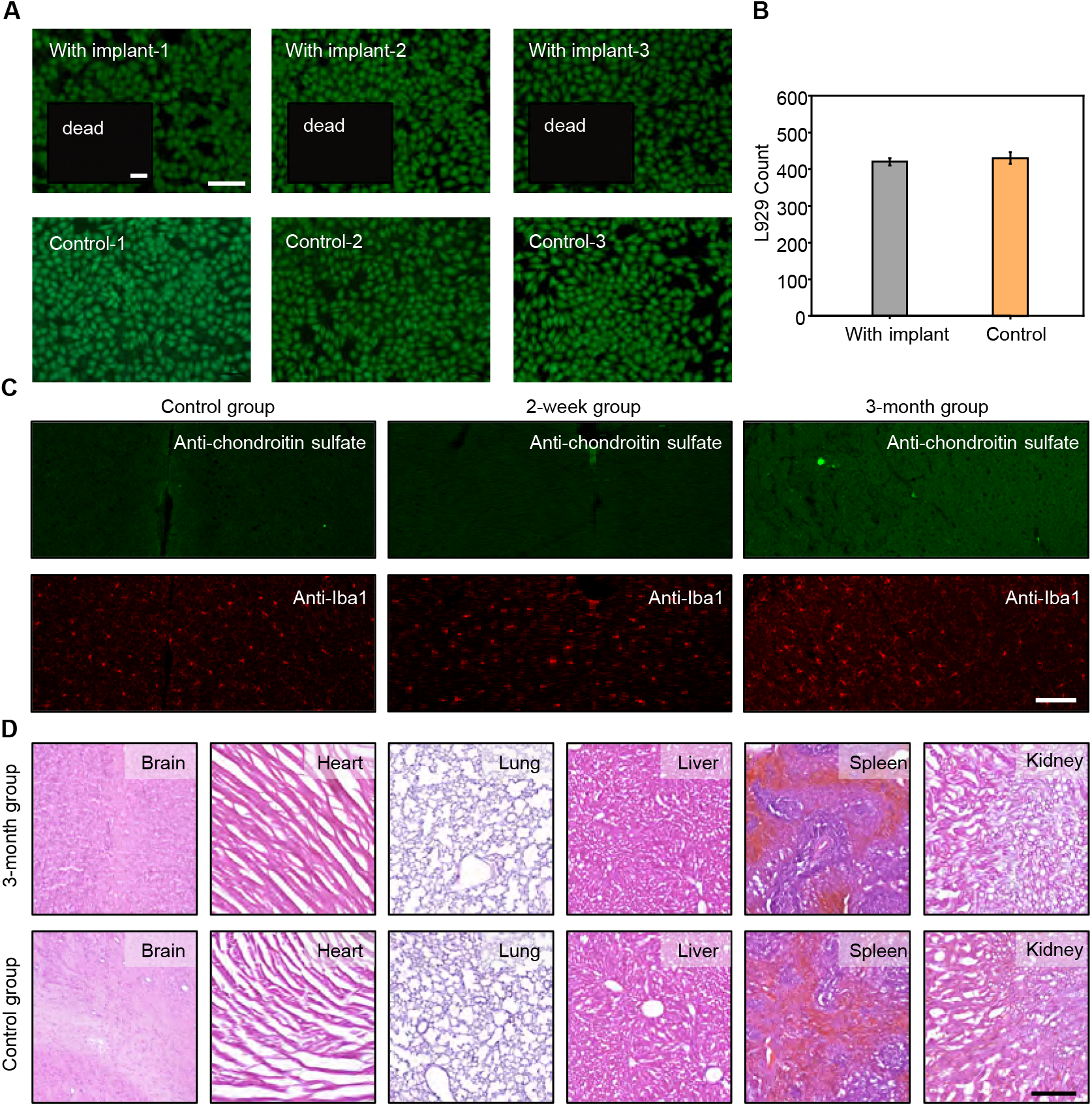
Biocompatibility of the magnetic implant. (**A**) Fluorescent images of fibroblast cells cultured on the magnetic implant for 2 weeks with Calcein-AM/propidium iodide. Green color (Calcein-AM) indicates live cells, and red color (propidium iodide) in insets indicates dead cells. Scale bars, 5 μm. (**B**) Comparison between control group and magnetic implants in cell proliferation. Error bars correspond to the calculated standard deviation from three measurements. (**C**) Immunofluorescent staining results (blue: cell nucleus; green: mouse anti-chondroitin sulfate; red: rabbit anti-Iba1) conducted on the brains of rats 2 weeks after implantation. Scale bar, 200 μm. (**D**) H&E-staining images of the brains, hearts, lungs, livers, spleens, and kidneys of rats with magnetic implants for 3 months. Scale bar, 200 μm.

During the mounting of the wearable device and the insertion of the implant, the relative position between them may influence the actuation and vibration of the micro-magnet, thereby affecting the accuracy in the measurement of pressure and other parameters. One approach to address this issue is to use the four Hall sensors in the fully integrated wearable device for positioning the magnetic implant (fig. S28). The accuracy reaches to 0.027 mm, 0.077 mm, 0.006 mm, 0.11° and 0.13° for x, y, z, θ and φ, respectively. Such small positioning errors permit accurate sensing (fig. S29, A to E). As a comparison, a shift of 0.1 mm along Y axis only results to an error of 0.1 mmHg (Δf/f0=0.032%) in pressure sensing, and a rotation of 0.1° around θ leads to an error of 0.07 mmHg (Δf/f0=0.023%, fig. S29F and text S3). An alternative means to eliminate the influence caused by changes of relative positions is to collect vibration waveforms of the magnetic implant at different relative positions for systematic calibration. The deep learning model yields a an R2 of 98.7% and a mean absolute error of 0.2 mmHg. The systematic calibration and deep learning model provide effective means to accurate sensing when the magnetic implant and wearable device locate at various relative positions. This is especially useful for in vivo applications, because the motion from the subject may change the relative positions of the wearable device and the magnetic implant. (figs. S30 to S33 and text S4).

We apply these implants to a rat brain and demonstrate the capability in wireless, continuous monitoring of CSF viscosity, ICP, and CSF glucose levels, as these parameters are important indicators for the diagnosis and treatment of traumatic brain injury (TBI), stroke, intracranial hemorrhage, acute meningitis, and brain inflammation (50–54). During craniotomy, dental cement (Kerr, Dyad Flow) securely attaches the magnetic implant to the skull with a drilled hole (diameter: 5.5 mm). The micro-magnet on the membrane faces towards the CSF through the craniotomy site (Fig. 4A). The computed tomography (CT) images in Fig. 4B and fig. S34 clearly visualizes of the location of the magnetic implant in the rat’s brain. After suturing and cleaning the wound, the rat recovers from anesthesia and can behave normally (Fig. 4, C and D and fig. S35). Placing the fully integrated wearable device on the rat’s head enables subsequent monitoring of physical and chemical parameters in the brain (Fig. 4, E and F).

**Fig. 4.**
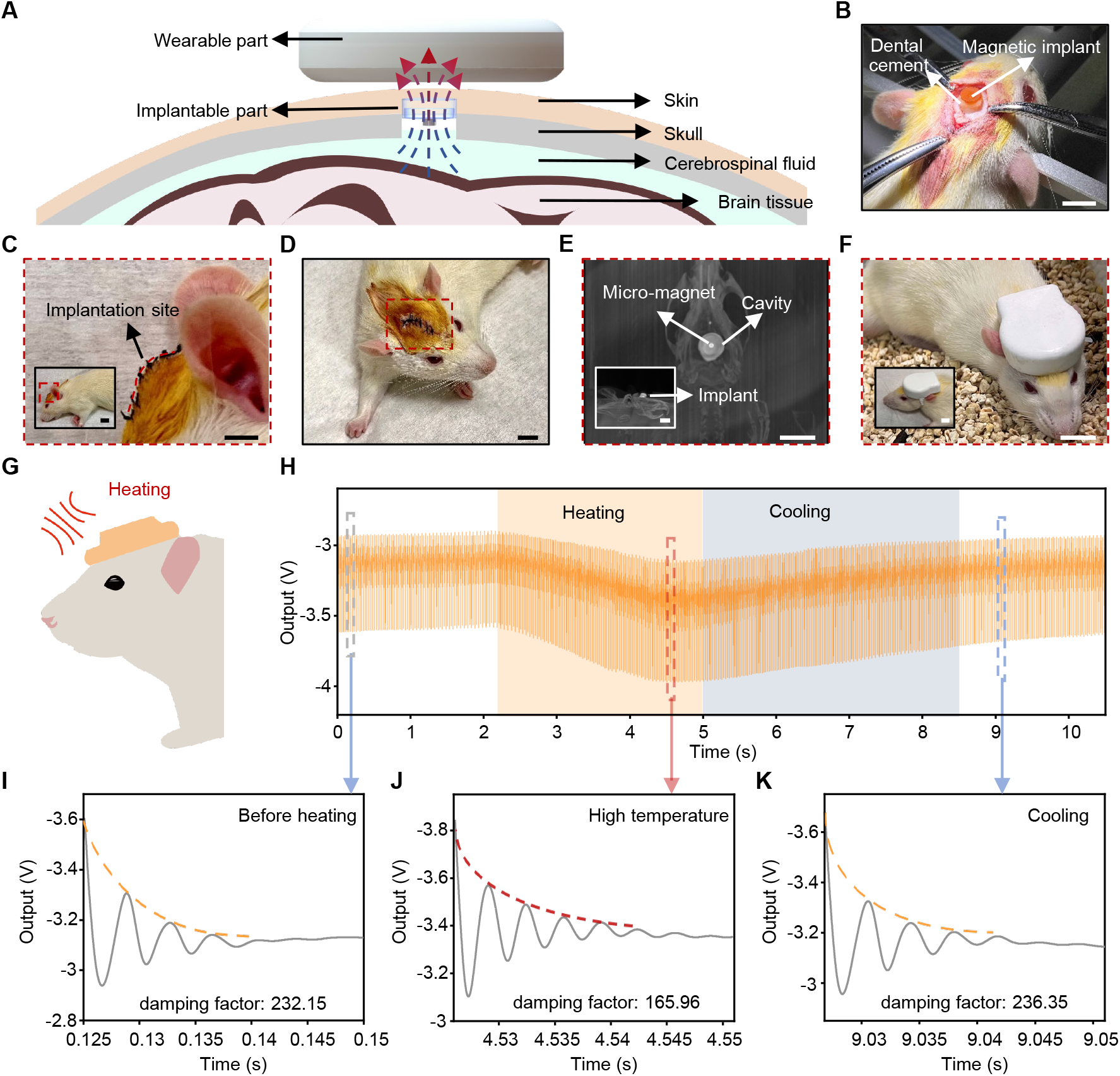
In vivo wireless sensing of CSF viscosity from the damping factor in rat’s brain. (**A**) Schematic illustration of the in vivo wireless measurement in rat’s brain. (**B**) CT image (top view) of the magnetic implant in the brain of a rat. The inset shows the side view CT image. Scale bars, 1 cm. (**C**) Optical image of a rat with fully sutured scalp after surgery. The inset is an optical image with a larger field of view. Scale bars, 5 mm. (**D**) Optical image of a freely moving rat after surgery. Scale bar, 5 mm. (**E**) CT image (top view) of the magnetic implant in the brain of a rat. The inset shows the side view CT image. Scale bars, 1 cm. (**F**) Optical image of a rat with a mounted wearable device. Scale bar, 1 cm. (**G**) Schematic diagram of altering CSF viscosity through heating. (**H**) Waveform captured from the wearable device during the actuation and damped vibration of the magnetic implant under different temperatures. (**I** to **K**) Magnified views of the damped vibration before heating (I), during heating (J), and after cooling (K).

In vivo adjustment of viscosity exploits the fact that higher temperature leads to lower viscosity of the fluid (55). Here, a hot air gun (100 °C) placed 6 cm away from the head of a rat increases the viscosity of CSF to ∼45 °C, thereby effectively reducing the viscosity of the fluid (56) (Fig. 4G). The wearable device mounted on the head of the rat then actuates the magnetic implant and records the intensity of the alternate magnetic field through the integrated TMR sensor. The waveform in Fig. 4H reveals that vibrations at high temperatures (i.e. lower viscosities) exhibit a noticeable slower damping compared to those at low temperatures (i.e. higher viscosities). The fitted damping factor is 232.15 before heating, gradually decreases to 165.96 during heating, and then returns to 236.35 after further cooling for ∼4 s. (Fig. 4, I to K).

We carry out in vivo measurement of ICP in a rat model and induce variations in ICP through abdominal compression (10) (Fig. 5A). The experiment exploits a commercial pressure sensor (JR-Intellicom, JR-3100D) inserted into the brain through the craniotomy, and a magnetic implant placed on top of a drilled hole of the skull (Fig. 5B and fig. S36). A close resemblance between the vibration frequency of the magnetic implant and the output of the commercial sensor indicates the accuracy and reliability of the magnetic implants in measuring ICP (Fig. 5C, implants: orange dots, commercial sensor: grey curve). Additional abdominal compression with various intensities (i.e. heavy, moderate, and light) result in different vibration frequencies of the magnetic implant. As shown in Fig. 5D and fig. S37, heavier abdominal compression leads to higher vibration frequency, indicating an elevated ICP. The magnetic implant maintains a consistent level of stability after implantation. Experiments performed on day 1 and day 3 both show obvious variations of vibration frequency during abdominal compression (Fig. 5E and fig. S38).

**Fig. 5.**
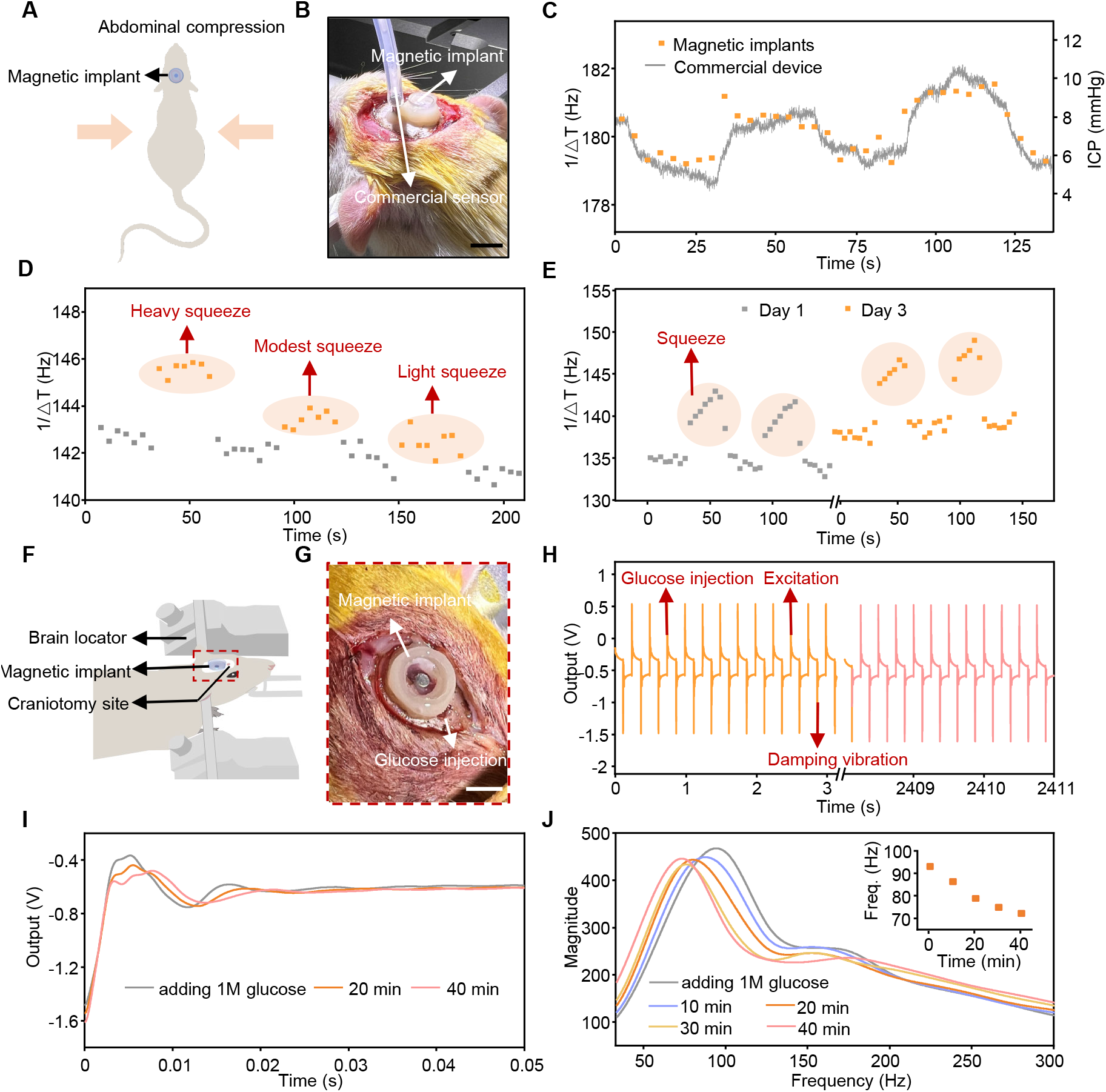
In vivo wireless sensing of ICP and CSF glucose level from the vibration frequency. (**A**) Schematic illustration of abdominal compression to adjust ICP. (**B**) Optical image showing the simultaneous measurement of ICP using the magnetic implant and a commercial pressure sensor. Scale bar, 1 cm. (**C**) Fluctuations of ICP captured from the magnetic implant (orange dots) and the commercial pressure sensor (grey curve) during abdominal compression. (**D**) Response of the magnetic implant under abdominal compression with various intensities. (**E**) Response of the magnetic implant during abdominal compressing on day 1 and day 3. Data points inside the light orange ellipses correspond to the ICP during squeezing. (**F** and **G**) Schematic illustration (**f**) and optical image (**g**) of the experimental setup for altering intracranial glucose concentration in rats. Scale bar, 5 mm. (**H**) Waveform captured from the wearable device during the process of glucose diffusion in CSF. (I) Magnified views of the one-cycle vibration waveforms over time after glucose injection. (**J**) FFT results of the vibration of the magnetic implant under different glucose levels. The inset shows the peak frequency in FFT as a function of time, indicating the diffusion of glucose across the CSF over time.

As illustrated in Fig. 5 (F and G), in vivo measurement of biochemicals, such as glucose concentration in CSF, demands an additional craniotomy site (diameter: 1 mm) on the skull located above the implantation site of the open-cavity magnetic implant. Injection of glucose solution through this site enables adjustment of glucose level in the CSF. Here, the injected solution with a small volume (100 μL) and a high concentration (1 M) can diffuse glucose into the CSF. The Con-A-modified MWCNTs on the micro-magnet surface gradually absorbs the glucose in the CSF, thereby increasing the mass of the membrane and the detected vibration frequency (Fig. 5, H and I). The processes of diffusion and absorption cover a period of ∼40 min, during which the vibration frequency of the magnetic implant decreases from 93.83 Hz to 72.88 Hz (Fig. 5J).

## Discussion

The chip-less, battery-less magnetic implants reported here can pair with a fully integrated wearable device to enable wireless monitoring of biophysical and biochemical parameters within the body, such as viscosity, pressure and glucose concentration. This system (i.e. the magnetic implant paired with the wearable device) offers qualitatively differentiated capabilities compared with existing wireless implantable sensors. First, the wearable device establishes a bidirectional interaction with the implant through magnetic field. This mechanism eliminates commercial chips, batteries and/or coils in the implants and does not require bulky readout equipment, thereby reducing the overall dimensions of the system for continuous monitoring to underpin pervasive healthcare. Second, the implants incorporate soft materials to generate vibration amplitudes two orders of magnitude larger than those of conventional MEMS, with tailorable vibration frequencies. Such designs facilitate the measurements in unshielded environment, and provide possibilities in multiplexed sensing using different frequency bands. Third, the vibration features of the magnetic implants after surface modification not only reflect their surrounding physical conditions, but also involve information of the concentration of a specific chemical. A deep learning model correlates the time-domain and frequency-domain data with various physical and chemical parameters, and makes the measurement insensitive to the relative positions between the magnetic implant and the wearable device. Fourth, the chip-less and battery-less nature of the implant promotes long-term biosafety, thereby supporting in vivo applications. Experiments in rat models validate the capabilities in wireless monitoring of CSF viscosity, ICP, and CSF glucose levels, offering personalized and customized references for the treatment of brain diseases, and the assessment of brain function and injuries. These advancements highlight our system as a complementary technology to existing wireless sensing modalities, serving as a unique tool for quantitative, continuous monitoring of a diverse set of biophysical and biochemical conditions.

Although the in vivo applications demonstrated here focus on the brain, the wireless sensing system can serve as an efficient diagnostic tool for many other regions inside the body in diverse application scenarios, including blood pressure and viscosity in cardiovascular system, contact force in dentistry and orthopedics, pressure in abdominal region, as well as the distributions and concentrations of proteins, peptides, small molecules, and cells in various regions inside the human body. Monitoring of these conditions outside of hospital settings is useful for the diagnosis, treatment, and management of a variety of acute and chronic diseases, such as traumatic injuries, heart failure, diabetes, cancer, liver cirrhosis, ascites and others. Many opportunities lie in the development of wireless biosensors by incorporating functional surface modifications (e.g. antibody, aptamer) to the magnetic implant. The resulting miniaturized system will allow for continuous sensing of a broader range of chemicals and biomolecules, all in a wireless manner. Additionally, downsizing the magnetic implants for minimally invasive insertion, increasing the distance of wireless actuation and recording to target at deep tissues, and leveraging the frequency band for multiplexed, multimodal sensing represents other important areas for future developments. These advancements will enhance the relevance and impact of our system in healthcare applications, and may renovate the diagnosis and treatment of a series of diseases.

## MATERIALS AND METHODS

### Circuit design of the wearable device

The wearable device comprised a high-sensitivity TMR sensor (TMR9002CP, DowayTech), a Bluetooth SoC (nrF52832, Nordic), a copper coil (1000 turns, Chuangli), an excitation circuit, four Hall sensors (BMM150, Bosch), and a lithium-ion battery (1200 mA, JieXun). The Bluetooth module, copper coil, and other components were powered directly by the lithium-ion battery with a voltage of 7.4 V. The excitation circuit utilized a DC-DC converter (TPS61170DRVR, Texas Instruments) to boost the voltage to 15 V. The Bluetooth SoC controlled the optocoupler switch (TLP187, Toshiba) within the excitation circuit, generating a square wave voltage input to the coil with a frequency of 4 Hz and a peak value of 15 V. This alternate voltage generated a square wave magnetic field that periodically attracted and released the magnet. The four Hall sensors estimated the initial position of the magnet on the membrane, while the TMR sensor captured the vibration signal of the micro-magnet. The analog output from the TMR sensor was then digitized by ADC and transmitted to a mobile terminal through Bluetooth. Subsequent signal processing extracted information related to the liquid viscosity, pressure, and glucose concentration.

### Fabrication of the PDMS cavity

Fabrication of the PDMS (Sylgard 184, Dow Corning) cavity started with mixing of the base and curing agent of PDMS at a weight ratio of 10:1. The sealed cavity was a tri-layer structure in PDMS, including a bottom substrate (thickness: 1 mm), a middle ring (thickness: 1 mm), and a top circular membrane (thickness: 40 μm). The open cavity in PDMS was a bi-layer structure, including a middle ring layer (thickness: 1 mm) and a top circular membrane (thickness: 40 μm). Laser-cutting defined the patterns of each layer, and uncured PDMS bonded three layers together to form a sealed cavity after heating at 80 °C for 40 min. A cylindrical magnet (D1001A-10, SuperMagneticMan) in neodymium iron boron with a diameter of 1.5 mm and a height of 0.8 mm coated with parylene-C layer (thickness: 10 μm) promoted the biocompatibility of the magnetic implants. Uncured PDMS enabled the bonding of the magnet to the center of the top membrane layer. Heating at 80 °C for 40 min cured the PDMS and completed the process.

### Fabrication of the vertical dense array of MWCNTs

The fabrication process involved growing a SiO_2_ buffer layer (thickness: 1 μm) on the Si substrate through thermal oxidation. This buffer layer prevented the direct fusion of the metal catalyst to the Si substrate. Following that, a thin layer of Fe (thickness: 2 nm), deposited through electron beam evaporation, served as the catalyst layer for MWCNTs growth. A quartz boat held the sample inside a reaction chamber, pumped to a pressure below ∼1.5 × 10^−2^ Torr. After introducing and stabilizing argon (Ar) gas at a flow rate of 7 sccm and hydrogen (H_2_) gas at a flow rate of 3 sccm, the pressure inside the chamber settled to be ∼0.36 Torr. The heating zone’s temperature gradually increased to 700 °C at a rate of 9.5 °C/min. Upon reaching a temperature of 695 °C, RF power supply switched on, setting the actual power to 250 W. After achieving a normal glow discharge inside the quartz tube, methane at a flow rate of approximately 14.7 sccm entered the chamber till the chamber pressure reached around 0.936 Torr, which initiated the growth of MWCNTs. The height of the MWCNTs had a positive relationship to the duration of the growth process. Once the designated growth process was over, RF power supply switched off. The flow of methane and hydrogen stopped and the equipment started to cooling down. A flow of 3.5 sccm of argon maintained to expedite the cooling process. When the temperature dropped to around 100 °C, the gas flow all stopped terminating the growth process.

### Transfer of MWCNTs from Si wafer to PDMS

The sample of MWCNTs (height: 50 μm) was on a Si wafer, with a thin layer of Fe (thickness: 2 nm) in between. The fabrication started with the spin-coating and curing process (80 °C for 40 min) of the first PDMS layer (thickness: 20.0 μm) on a polyethylene terephthalate (PET) film. Then, the uncured second PDMS layer (thickness: 20 μm) spin-coated on the first PDMS layer trapped the MWCNTs on the wafer. After curing the second PDMS layer at 80 °C for 40 min, it firmly held the MWCNTs to allow for the release from Si wafer. Laser cutting patterned the PDMS membrane with MWCNTs into circular shape with a diameter of 1.5 mm.

### Modification of MWCNTs with Con-A

The modification began with immersing the membrane in a 3 mg/ml 1-pyrenebutanoic succinimidyl ester (PBSE, Xianding Bio-Technology Co., Ltd) solution in N,N-Dimethylformamide (DMF, CONCORD Technology) for 4 h. Then, immersing the membrane with DMF and phosphate buffer solution (PBS), and soaking it in PBS (SinoDetech) containing 1 mM calcium ions, 1 mM manganese ions, and 3 mg/ml Con-A (Shanghai Yuanye Bio-Technology Co., Ltd) for 30 min completed the binding process. After washing away the excess Con-A with PBS, immersing in a 0.1% ethanolamine (Energy Chemical) aqueous solution for 1 h blocked unbound Con-A binding sites. Subsequently, immersing the sample in a 0.1% Tween-20 (Coolaber) solution for 30 min blocked exposed MWCNTs and reduced non-specific binding. Finally, washing the membrane with PBS completed the specific modification of MWCNTs for the selective binding with glucose molecules. The resulting membrane was suitable for glucose sensing applications.

### Viscosity testing environment

Different ratios of glycerol-water solutions had distinct viscosities. During the test, the magnetic implants located in the liquid environment with various viscosities. The wearable device positioned above the magnetic implants collected the signals resulting from changes in the magnetic field caused by the vibration of the micro-magnet. The damping of the vibration depended on the liquid viscosity.

### Pressure testing environment

The pressure at the lower end of the customized U-tube had a positive relationship with the liquid (artificial CSF, MREDA) height difference between the two pipes. Positioned above the magnetic implant at the lower end, the wearable device collected the signals resulting from changes in the magnetic field caused by the vibration of the micro-magnet. The vibration frequency depended on the liquid pressure.

### Glucose testing environment

The MWCNTs-PDMS membrane, modified with Con-A, adhered to the micro-magnet using uncured Ecoflex Gel. Curing at room temperature for 10 min promoted the adhesion between the MWCNTs-PDMS membrane and the micro-magnet. Then, fixing the open-cavity actuators with MWCNTs-PDMS membrane into the liquid environment (artificial CSF) with different concentrations of glucose, the wearable device positioned below the open-cavity actuators can collect the signals resulting from changes in the magnetic field caused by the vibration of the magnet. The vibration frequency depended on the concentration of glucose.

### Automated calibration at different relative positions

During calibration, the magnetic implants located in liquid environment (artificial CSF) with a fixed position. A customized 4-axis automated displacement platform held the wearable device. The platform moved along X, Y, Z axes and rotated around X axis to adjust the relative position between the wearable device and the magnetic implant. In each position, the wearable device actuated the magnetic implant and recorded the intensity of the alternate magnetic field. The recorded signals served as training samples in deep learning.

### Deep learning model

A PowerLab computer interface (Model 16/35, ADInstruments) collected the signal at a frequency of 200 kHz. A peak detection algorithm enabled extraction of the vibration signal. The time domain and frequency domain data served as features for deep learning. The actual values of viscosity, pressure, and glucose concentration were truth values in the deep learning model. The vibration signals for calibration of viscosity, pressure, and glucose concentration in benchtop test were training samples. The preprocessed signal from the in vivo test were testing samples. The MLP regression algorithm allowed for reliable regression of the variables (e.g. viscosity, pressure, and glucose concentration) from the testing samples.

### In vitro cytotoxicity evaluation

The proliferation experiment began with soaking the magnetic implants in 75% alcohol, disinfecting it with ultraviolet light irradiation and washing it with PBS for three times. Then, preparation of L929 cells involved seeding the cells in 96-well plates at a density of 5000 per well along with the magnetic implants. The control group used RPMI-1640 basal medium with 10% fetal bovine serum (FBS). After 3 days ofincubation, Calcein AM/propidium iodide (Biyuntian Co,Ltd.) staining was carried out, with green(Calcein-AM) for live cells and red (propidium iodide) for dead cells. The fluorescence images were obtained with fluorescence microscopy (Axi VertAZeiss).

### Surgical procedures

All procedures associated with animal studies followed recommendations with the ethical guidelines of the Institutional Animal Center Peking University. The Institutional Animal Care and Use Committee (IACUC) at Peking University (Protocol No. FT-HanMD-1) approved the protocol. Male rats (SD, age at initiation of the treatment: at least 9 weeks, but no more than 15 weeks, purchased from Beijing Vitalstar Biotechnology Co., Ltd.) were acclimated up to 5 days before surgery. Animals were anaesthetized using isoflurane gas during the implantation surgery and measurement. Surgical procedures removed the furs on the head and created a craniotomy (5.5 mm diameter) in the skull located in the parietal lobe area. Next, we carefully cut the dura mater and introduced saline into the craniotomy to establish a liquid environment for testing purposes. After cleaning the implantation site and the surface of the magnetic implant, dental cement enabled bonding of the magnetic implant to the designated site of the skull. Disinfecting and suturing the wound completed the implantation surgery, followed by administering penicillin for anti-inflammatory effects and injection of meloxicam for pain relief. Closing the scalp completely sealed the magnetic implant inside, without any transcutaneous wires. Placing the wearable device on the head of the rat enabled wireless actuation of the magnetic implant and wireless recording of the dynamic magnetic field.

### In vivo biocompatibility test

All procedures associated with animal studies followed recommendations with the ethical guidelines of the Institutional Animal Center Peking University. The Institutional Animal Care and Use Committee (IACUC) at Peking University (Protocol No. FT-HanMD-1) approved the protocol. After the 2-week and 3-month monitoring window, rats implanted with magnetic implants (2 weeks: n=4; 3 months: n=2) and a rat without implantation (control group, n=1) were dissected through a surgical operation, removing organs including brain, heart, lung, liver, spleen and kidney for histological investigations. Subsequently, organs went through 1-week fixing in polyoxymethylene and 5-week dehydration in sucrose solution. Then, the sliced samples of the brain tissue were stained by rabbit anti-Iba1 (1:500, Abcam) to label microglia, mouse anti-chondroitin sulfate (1:400, Sigma) to detect production of chondroitin sulfate proteoglycan. In the meanwhile, organs including brain, heart, lung, liver, spleen and kidney went through hematoxylin and eosin (H&E) staining. After staining, all the samples were examined under a digital slide scanning system (Pannoramic Scan, 3Dhistech).

### In vivo biophysical and biochemical sensing on rat models

The experiments were conducted in accordance with the ethical guidelines of the Institutional Animal Center Peking University and with the approval of the Institutional Animal Care and Use Committee of the Peking University in Beijing (Protocol No. FT-HanMD-1). Localized heating to the brain through a hot air gun (OLT8586, Oulitai) altered the viscosity of the CSF. Abdominal compression applied gentle pressure to the abdomen of the rat, resulting to a change of ICP. To adjust the glucose concentration in the CSF, the procedure introduced a high-concentration glucose solution (100 μL, 1 M) into the craniotomy site. The frequency and damping rate of the vibration signals provided information related to CSF viscosity, ICP, and CSF glucose level. Additionally, a commercial pressure sensor (JR-3100D, JR-Intellicom) inserted into the brain from a flexible tube in polyethylene (diameter: 1 mm) allowed for calibration purposes. In all the in vivo experiments, dental cement sealed the hole on the skull and fixed the interface between the sensor and the skull to ensure secure and reliable connections.

## Data availability

The data that support the findings of this study are available from the corresponding author upon reasonable request.

## Code availability

The Code that support the findings of this study are available from the corresponding author upon reasonable request.

## Acknowledgements

We thank P. Cheng and H. Zhang for providing valuable suggestions. This work was supported by the National Natural Science Foundation of China (No. 62104009); the Emerging Engineering Interdisciplinary Project, Peking University, the Fundamental Research Funds for the Central Universities; and Peking Nanofab Laboratory.

## Author contributions

J.W. and M.H. conceived the idea and designed the research; Z.N. and J.W. designed the circuits in wearable device; J.W. and J.X. carried out the animal experiments with the assistance from S.Y. and W.M.; Z.Z., M.Z., and J.W. performed the surface modification experiment; S.Y. and J.W. conducted the biocompatibility test; J.W., Y.L. and J.W. developed the deep learning algorithm; J.W. conducted the benchtop test and calibration test with the assistance from Z.X., X.L., C.X., P.Z., Y.W., J.Z., Y.W., S.Z.; J.W. and M.H. wrote the manuscript, and all authors provided feedback.

## Competing interests

The authors declare no competing interests.

## Notes

### Competing Interest Statement

The authors have declared no competing interest.

