## Supplementary Materials for "Millimeter-scale magnetic implants paired with a fully integrated wearable device for wireless biophysical and biochemical sensing"

### 1. Supplementary Text

#### Text S1. Comparation between the magnetic implants and other wireless implantable devices

***(1) Bluetooth***

Compared to implantable devices that use Bluetooth chips for signal transmission, the magnetic implants introduced here do not require integrated silicon-based circuits, resulting in reduced rigidity and improved biocompatibility. The magnetic implants are passive, and do not rely on power supplied such as implantable battery or wireless power transmission. The elimination of battery and coils reduces the overall dimension of the magnetic implants to millimeter scale, whereas implantable wireless devices based on Bluetooth are usually in centimeter scale (*1*,*2*).

***(2) NFC and RF resonance***

Wireless implantable devices based on NFC or RF resonance are passive (*3*-*9*). The implantable part can be very small, but they require large reading coils and signal processing circuits that increase the dimension of the data acquisition part. The large data acquisition device hinders the wearable and portable applications of the system. Additionally, these implantable devices are more susceptible to the influence of intermediate mediums such as skin, tissue, bones, and relative displacement.

***(3) Ultrasound***

Ultrasound can deliver energy to implantable devices integrated with piezoelectric materials. The vibration of the piezoelectric material can also be captured by external ultrasound transducers. Neural dusts represent one example of using ultrasound for wireless sensing (*10*-*12*). However, the implantable part requires rigid silicon chips, and the external part demands ultrasound probes and associated circuits which are relatively large. In addition, the conduction of ultrasound is significantly affected in the presence of intermediate mediums like bones.

***(4) Light irradiation***

Light has limited penetration through human skin and is significantly influenced by the material and color of intermediate mediums (*13*,*15*). Although wide-field illumination in the second near-infrared spectral window (NIR-II) can penetrate the skull, the penetration depth is limited to ~6 mm and additional light sources and imaging equipment are required (*15*).

***(5) Other magnetic implantable devices***

Existing magnetic implants rely on large external coils or magnets for actuation (*16*-*18*). Sensing functions from these magnetic implants require large external coils or imaging equipment that are not suitable for wearable applications (*19*,*27*).

#### Text S2. Scaling law of pressure sensing

A series of FEA generates the scaling law of the vibration frequency:

$$\mathrm{f} = \mathrm{k}D^{-\frac{2}{3}}T^{\frac{1}{3}}Y^{\frac{1}{6}}P^{\frac{1}{3}}M^{\frac{1}{2}}$$

where $f$ is the vibration frequency, $k$ is a constant, $M$ is the mass of the micro-magnets, $P$ is the pressure that the surrounding liquid environment applied on the magnetic implant, $D$, $T$, and $Y$ represent the diameter, thickness, and Young's modulus of the elastic membrane, respectively.

In FEA, the ranges of these parameters are set to:

$T$ from 35 μm to 70 μm,

$D$ from 5.5 mm to 7.25 mm,

$Y$ from 300 kPa to 1000 kPa,

$P$ from 5 kPa to 55 kPa,

$M$ from 0.09g to 0.104 g.

These values fall within the range of parameters of the magnetic implants used in experiments. The scaling law captures the relationship between $f$, and $T$, $D$, $Y$, $P$ and $M$ in this range, with $R^2$>95% in linear regression in logarithmic coordinate.

#### Text S3. Measuring the relative position using four Hall sensors

We can measure the position of the magnetic implants by using the four Hall sensors integrated in the wearable device. To evaluate the accuracy in positioning, a customized 5-axis motor automatically alters the position and angle of the wearable device, while keeping the magnetic implant at a fixed position. Utilizing the 5-axis motor, we collect the outputs of four Hall sensors when the magnetic implant locates at different relative positions to form a training set (X axis covers a range of 8 mm with a step of 1 mm, Y axis covers a range of 9 mm with a step of 1 mm, Z axis covers a range of 3 mm with a step of 1 mm, θ covers a range of 3° with a step of 1°, φ covers a range of 3° with a step of 1°). Afterwards, exploiting the same setup, we move the magnetic implant at various relative positions, and use the outputs of the Hall sensors as test set. An MLP regression algorithm allows us to locate the magnetic implant with high accuracy (mean absolute error of 0.027 mm for X axis, 0.077 mm for Y axis, 0.006 mm for Z axis, 0.11° for θ and 0.13° for φ, fig. S29, B to F). Thus, by using the output from the four Hall sensors, we can adjust the mounting location of the wearable device to maintain a fixed relative position to the magnetic implant.

#### Text S4. Calibration at different relative positions

The actuation strength from the coil and the vibration waveform of the magnetic implant depend not only to the external pressure, but also to the relative positions between the magnetic implant and the wearable device. To mitigate the influence of relative position, we perform calibrations at various relative positions using the customized 4-axis motor (wearable device at different positions and angles, magnetic implant at a fixed position). The calibration generates a more comprehensive dataset for the deep learning model.

Specifically, the movements along X, Y and Z axes cover a range of 4 mm with a step of 1 mm, the rotational angle (θ) around X axis has a range of 3° with a step of 1°. The initial distance between the coil and micro-magnet is 2 mm. A moving range of 4 mm in Z axis means that the distance changes from 2 mm to 6 mm. After the distance reaches and exceeds 4 mm, the rotation from 0 to

3° initiates. At shorter distances, the space occupied by the device hinders the rotation at relatively large angles. The implant is positioned horizontally above the wearable device in the X-Y plane, and we systematically explore 25 different positions within a 4 by 4 mm area on the X-Y plane, with a 1 mm interval (fig. S31).

As we move along the X-axis, the TMR sensor capture a square wave waveform characterized by a stepped pattern (fig. S32, A and B). The variations between the steps correspond to the displacement of the implants along the X-axis. Movement of the wearable device along Y-axis also led to variations in the square wave (fig. S32C). Afterwards, the motor moves the wearable device along Z-axis (fig. S32D), with a range of 4 mm and a step of 1 mm. In each Z-axis position, we collect 25 vibration waveforms by moving the wearable device across the X-Y plane (i.e. moving along X-axis from 0 to 4 mm, moving along Y-axis from 0 to 4 mm). The diagram demonstrates that changes in the Z-axis position result in a decrease in the peak-to-peak value of the square wave signal (fig. S32E). Additionally, we explore the impact of the relative angle between the implants and the wearable device (fig. S32F). During the traversal of positions in the aforementioned three dimensions (i.e. X, Y and Z axes), we collect vibration data from various positions at four different angles (0°, 1°, 2° and 3°) when the distance between the implants and the wearable device is over 4 mm. As illustrated in the diagram, the deviation angle also influences the vibration waveform (fig. S32G). The above measurements produce vibration waveforms at 350 different relative positions (5 in X axis, 5 in Y axis, 5 in Z axis, 4 in rotation when $Z>2$ mm). In each position, the pressure changes from 0 mmHg to 12 mmHg with a step of 3 mmHg. These waveforms are collected and utilized as the training set for deep learning (fig. S33A).

This approach enables accurate sensing of pressure, regardless of the relative positions of the magnetic implant and the wearable device. The test set consists of 500 samples independent of the training set. Results in fig. S33B indicate an $R^2$ of 95% and a mean absolute error within 1 mmHg. In addition, reducing the moving steps calibration can further improve the accuracy. As a demonstration, we gather vibration waveforms from various positions with smaller moving steps (X, Y axes cover a range of 1 mm with a step of 0.2 mm, Z axis covers a range of 1 mm with a step of 0.25 mm, $\theta$ cover a range of 1° with a step of 0.2°). Utilizing them as the training dataset and another 500 independent samples as the test set, we achieve an $R^2$ of 99% and a mean absolute error of 0.2 mmHg (fig. S33C). This approach caters to many in vivo applications, where the deformations of the skin or organs alter the relative positions/angles of the wearable device and magnetic implants. Despite these changes, the wearable device can still effectively excite and capture the vibration of the micro-magnet, enabling the regression of the true value of the pressure with minimal error.

#### 2. Supplementary Figures

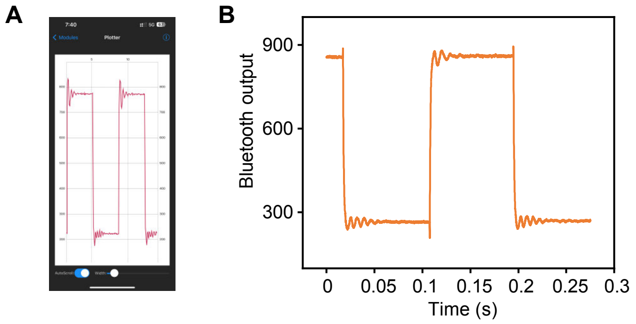

**Fig. S1. Bluetooth transmission of the output signal from TMR sensor.** (**A**) Screenshot of the signal transmitted to a mobile phone through Bluetooth. (**B**) Signal from the TMR sensor transmitted via Bluetooth.

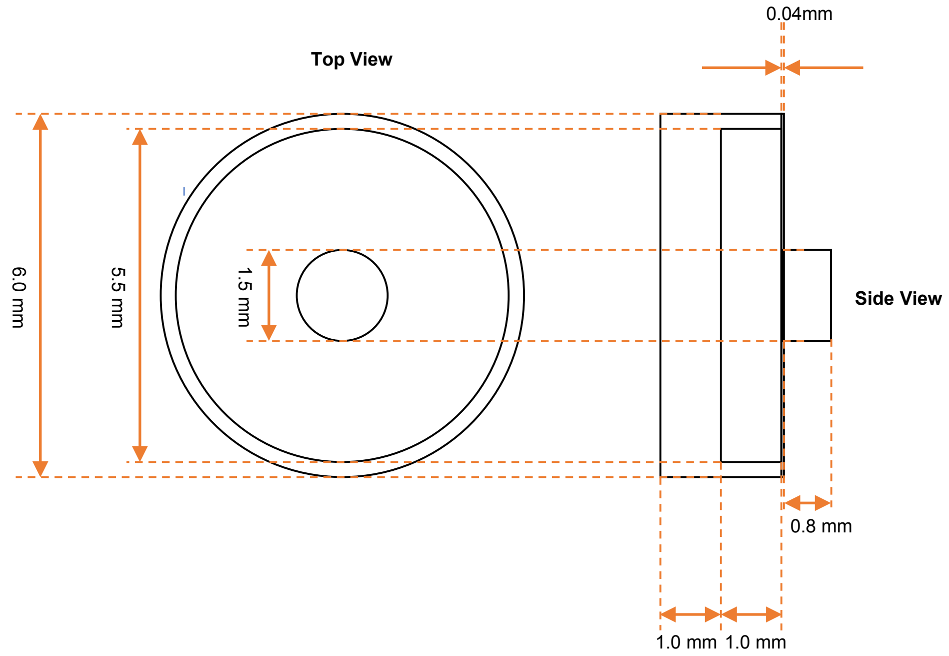

**Fig. S2. Dimensions of the magnetic implant with a sealed cavity.** Left: top view. Right: side view.

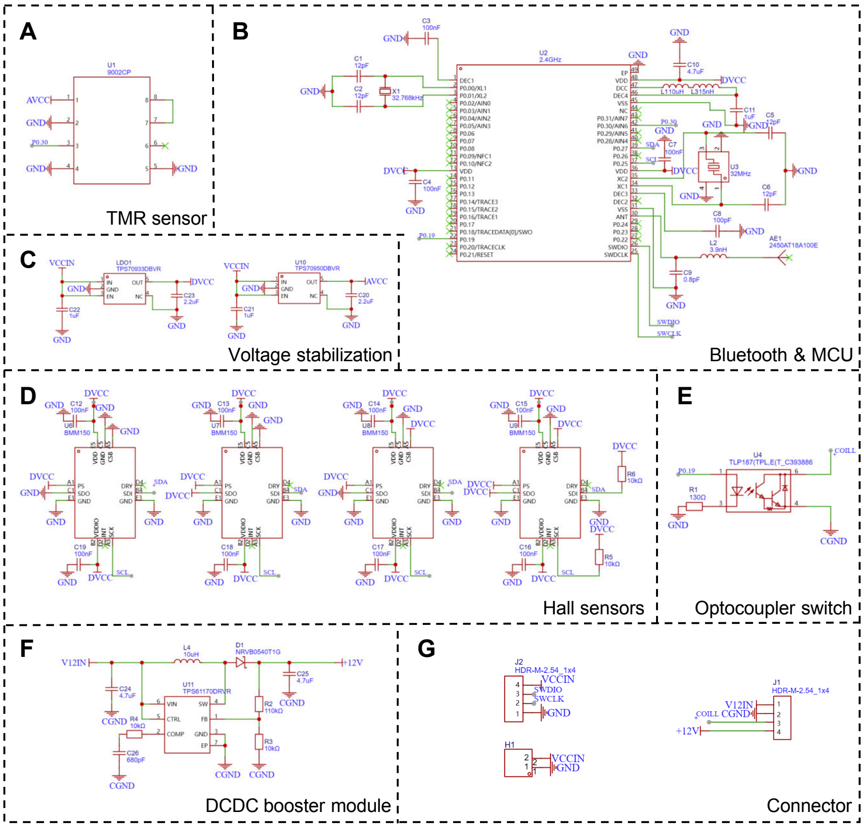

**Fig. S3. Circuit of the fully integrated wearable device.** (**A**) TMR sensor for wireless sensing. (**B**) Bluetooth and microcontroller unit (MCU) for wireless signal transmission. (**C**) Voltage stabilization unit. (**D**) Hall sensors for localization of the magnetic implant. (**E**) Connectors. (**F**) DC-DC booster module connected to the coil for actuation of the magnetic implant. (**G**) Optocoupler switch for generating a square wave voltage.

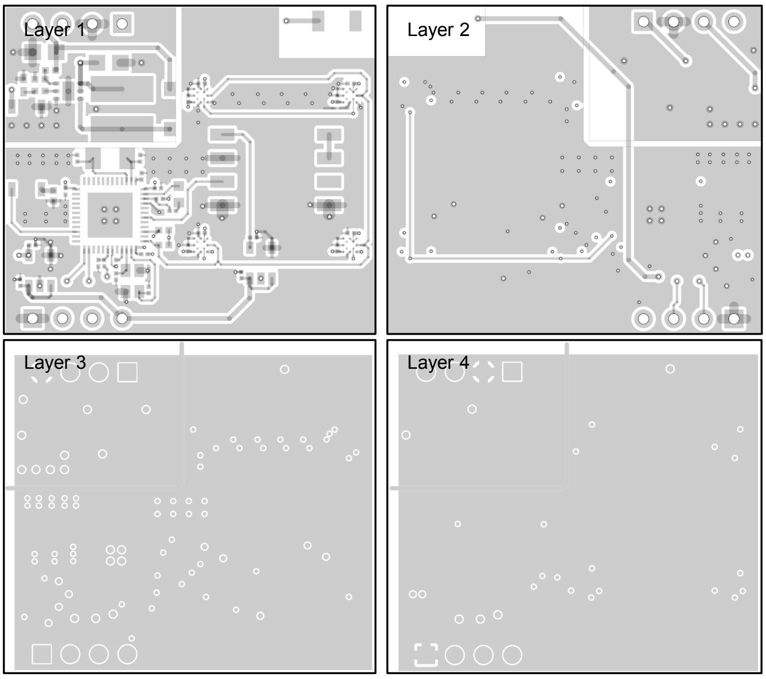

**Fig. S4. Layout designs of the printed circuit board (PCB) circuit for the fully integrated wearable device**.

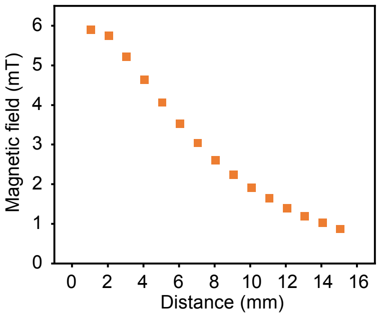

**Fig. S5. Magnetic field generated by the coil (input current: 100 mA) as a function of distance**.

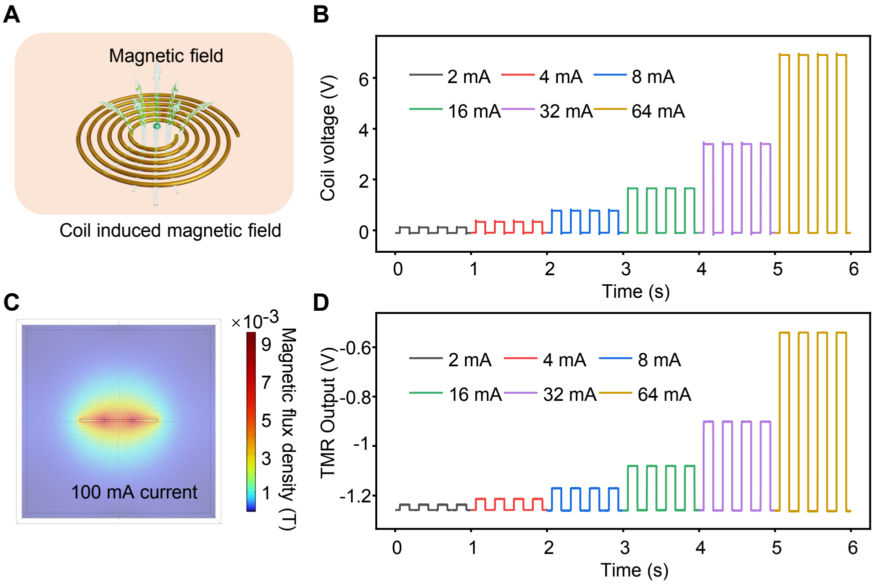

**Fig. S6. Coil for actuation of the magnetic implant.** (**A**) Schematic illustration of the coil for actuation. (**B**) Measured voltage from the coil under square wave input currents at different amplitudes. (**C**) FEA prediction of the magnetic field induced by the coil under a current of 100 mA. (**D**) Response of the TMR sensor under square wave input currents at different amplitudes.

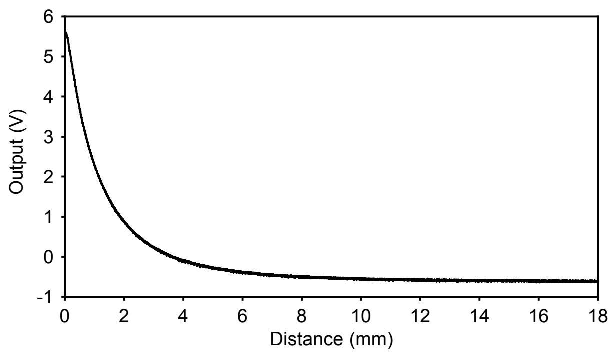

**Fig. S7. Magnetic field of a micro-magnet (diameter: 1.5 mm, height: 0.8 mm) detected by the TMR sensor at various distances.** The y axis is the output voltage from the TMR sensor.

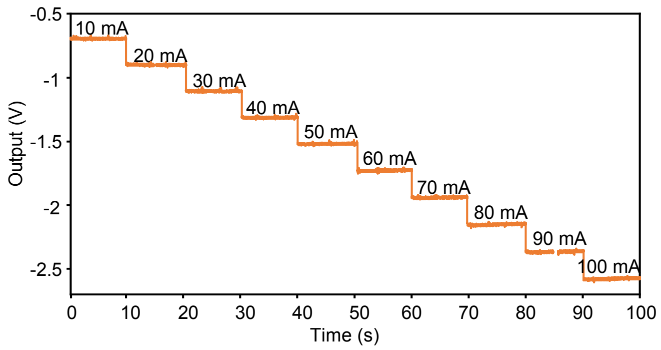

**Fig. S8. Magnetic field of a coil with different input currents detected by the TMR sensor.** The distance between the TMR sensor and the coil is 3 mm. The input current ranges from 10 mA to 100 mA.

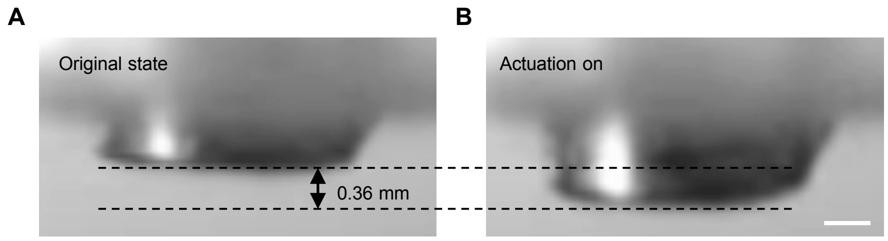

**Fig. S9. Optical images of the micro-magnet under excitation from the coil.** (**A**) Optical image of the micro-magnet at original position. (**B**) Optical image of the micro-magnet dragged by the coil's magnetic field. Input current: 100 mA. Distance between the coil and the magnet: 5 mm. Scale bar, 0.4 mm.

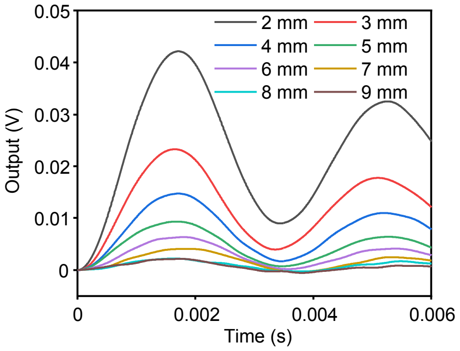

**Fig. S10. Vibration signals from the magnetic implant captured by the wearable device at varying distances.**

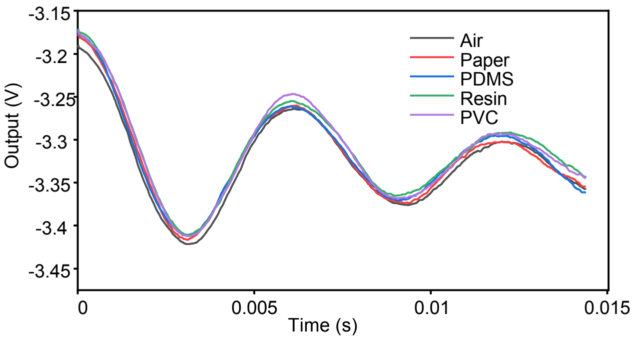

**Fig. S11. Vibration signals captured by the wearable device when inserting intermediate mediums between the magnetic implant and the wearable device.**

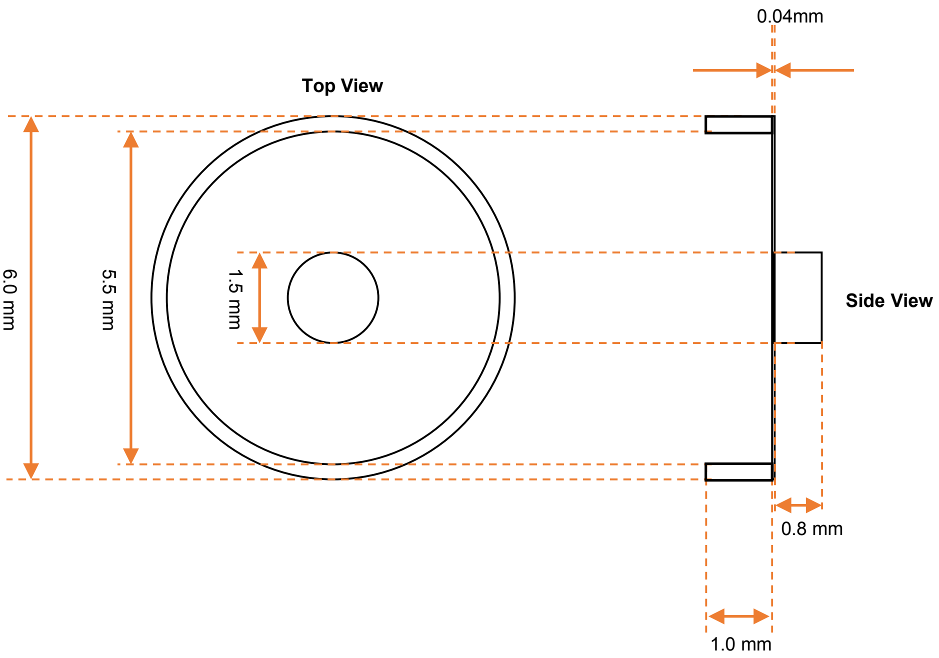

**Fig. S12. Dimensions of the magnetic implant with an open cavity.** Left: top view. Right: side view.

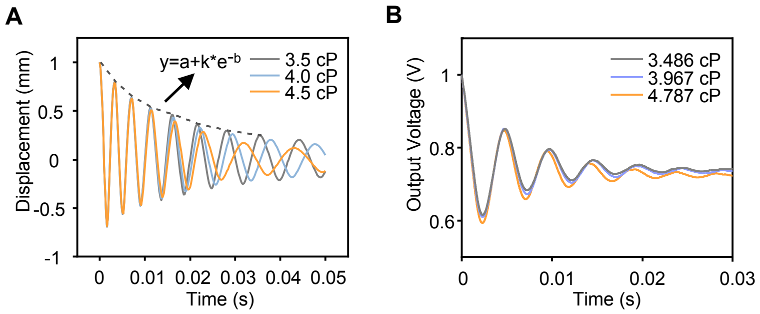

**Fig. S13. Response of the magnetic implant under different viscosities.** (**A** and **B**) FEA predictions (A) and experimental measurements (B) of the vibration signals from the magnetic implant (sealed cavity) under different viscosities.

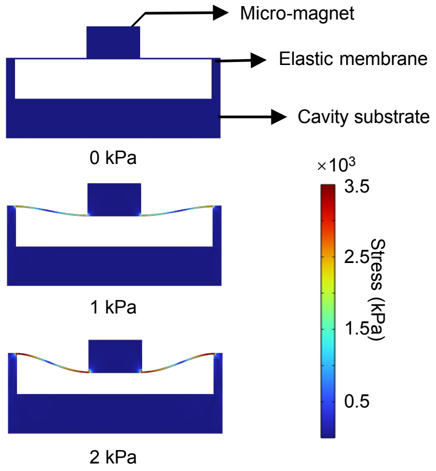

**Fig. S14. Deformation and stress distribution of the magnetic implant (sealed cavity) under different pressures.**

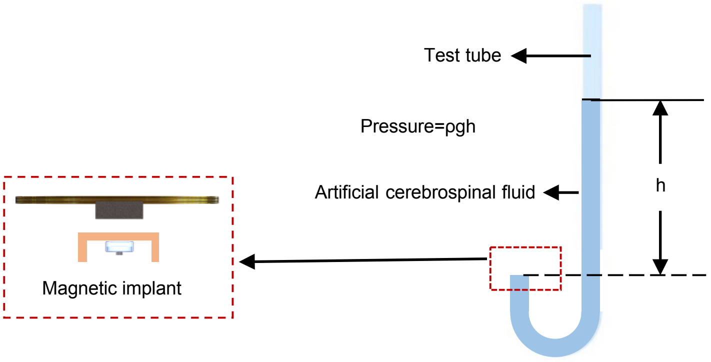

**Fig. S15. Schematic illustration of the pressure tests in liquid environment.**

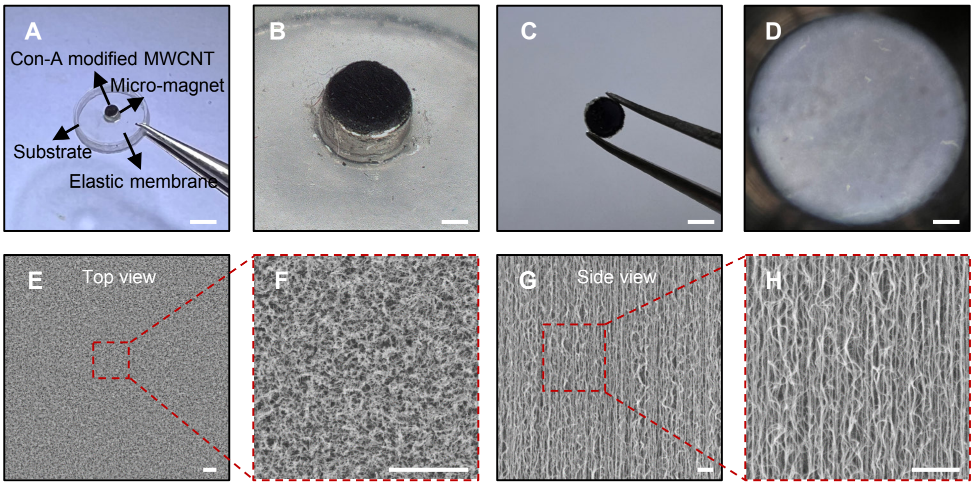

**Fig. S16. Magnetic implant with an open cavity for glucose sensing.** (**A**) Optical image of the magnetic implant with an open cavity. Scale bar, 3 mm. (**B**) Optical image of the MWCNTs on micro-magnet. Scale bar, 1 mm. (**C**) Optical image of the MWCNTs on PDMS/PET substrate. Scale bar, 1 mm. (**D**) Microscopic image of the MWCNTs. Scale bar, 100 μm. (**E** to **H**) Scanning electron microscope (SEM) image of the MWCNTs ((E) and (F): top view, (G) and (H): side view). Scale bars, 3 μm.

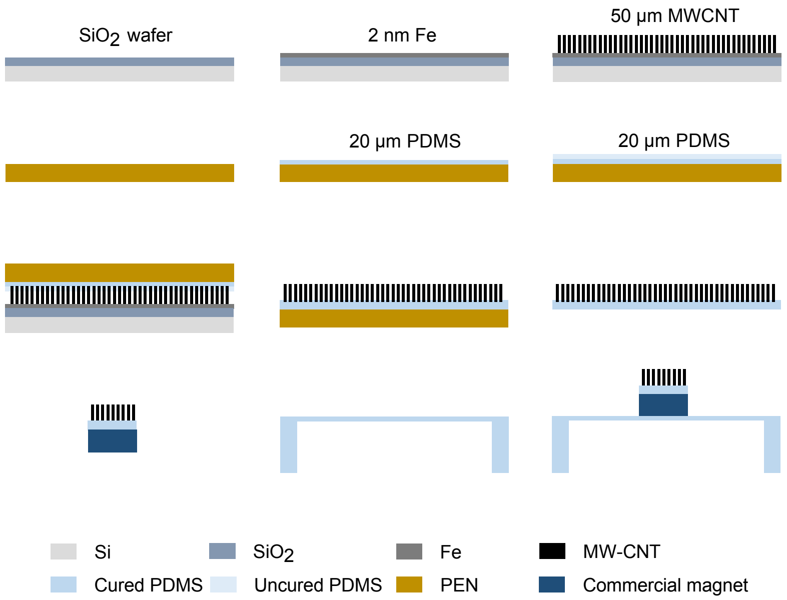

**Fig. S17. Schematic illustration of the fabrication process of the open cavity device with MWCNTs.**

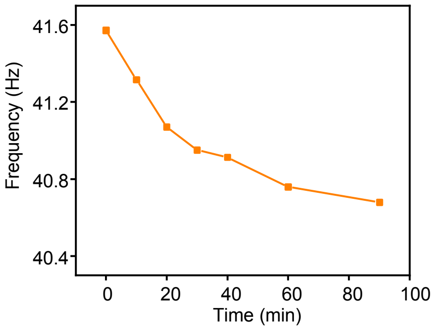

**Fig. S18. Frequency shift of the magnetic implant (open cavity, diameter: 8 mm) when adding 1 mL of 1 M glucose aqueous solution to 50 mL of artificial CSF.**

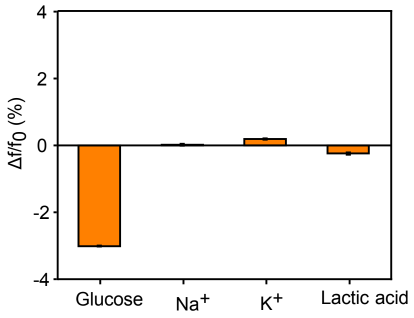

**Fig. S19. Selectivity of Con-A binding glucose against NaCl, KCl and Lactic acid.** Error bars correspond to the calculated standard deviation from five measurements.

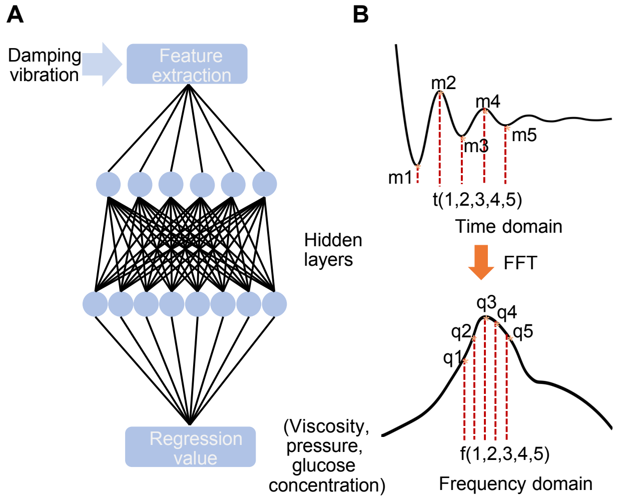

**Fig. S20. Deep learning regression algorithms.** (**A**) MLP regression model for biophysical and biochemical sensing. (**B**) Schematic illustration of the time-domain waveform and the frequency-domain waveform obtained through FFT**.**

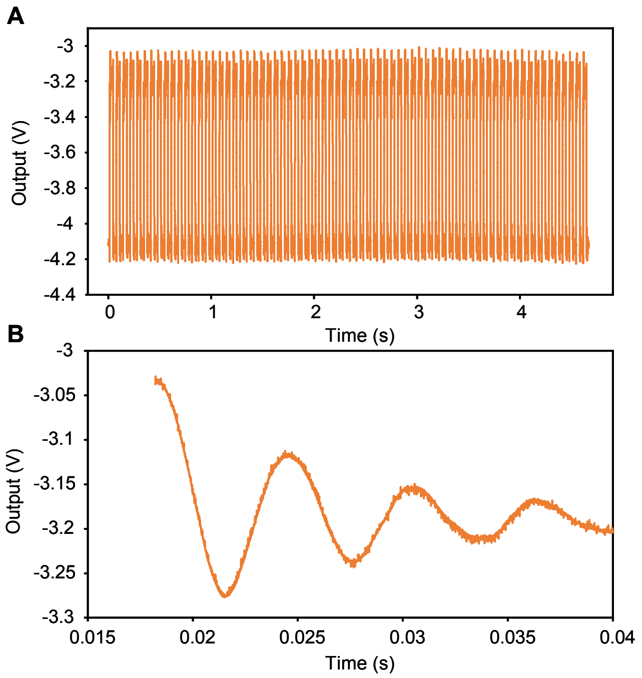

**Fig. S21. Coil excitation at a frequency of 15 Hz.** (**A**) Output of the TMR sensor under 15 Hz excitation. (**B**) Magnified view of a single cycle of damped vibration.

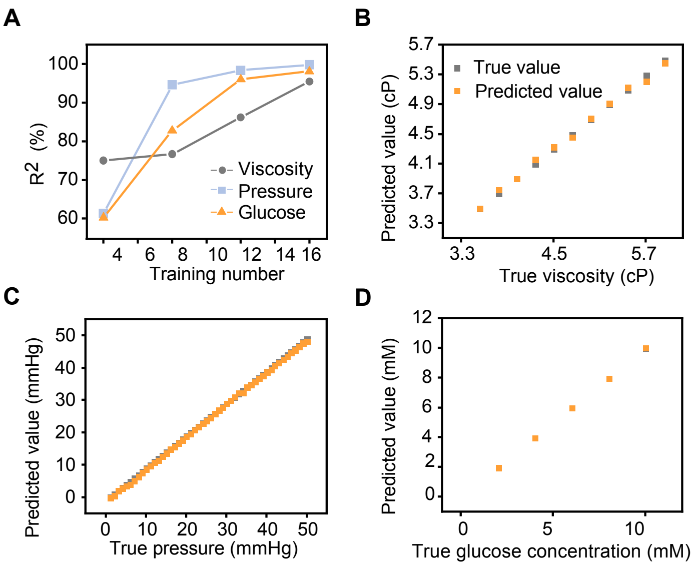

**Fig. S22. MLP regression results.** (A) Relationship between training number and the regression $R^2$. (B to D) Regression results for viscosity (B), pressure (C) and glucose concentration (D).

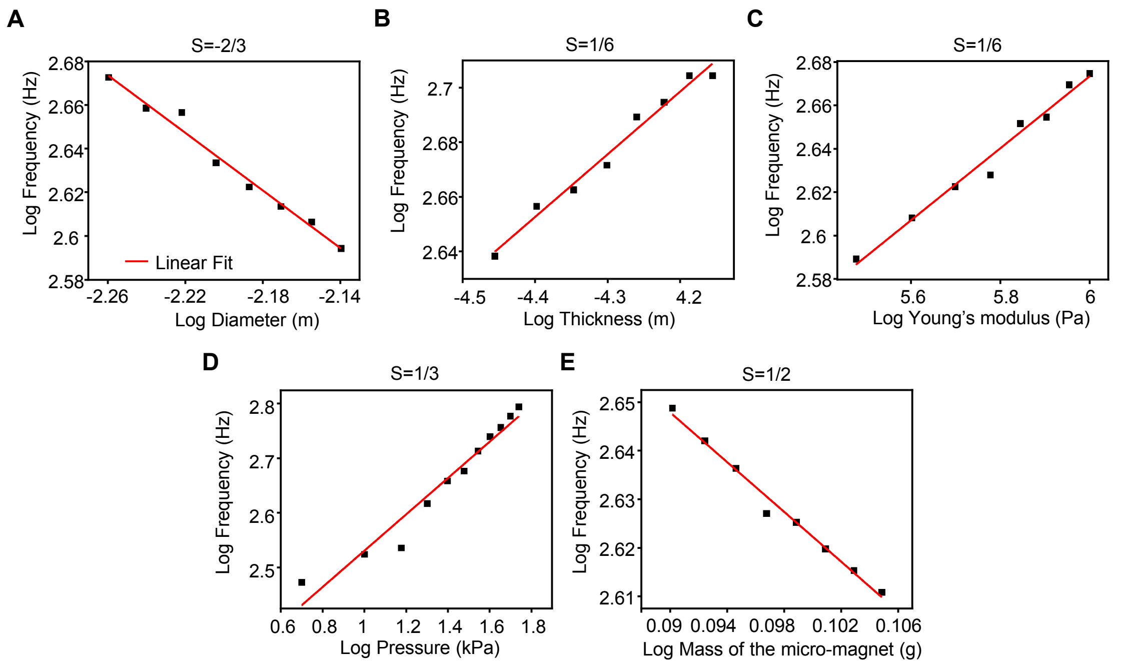

**Fig. S23. Scaling law of the magnetic implants with sealed cavity.** (**A**) Influence of the diameter of elastic membrane on vibration frequency. (**B**) Influence of the thickness of elastic membrane on vibration frequency. (**C**) Influence of the Young's modulus of elastic membrane on vibration frequency. (**D**) Influence of the external pressure on vibration frequency. (**E**) Influence of the mass of the micro-magnet on vibration frequency.

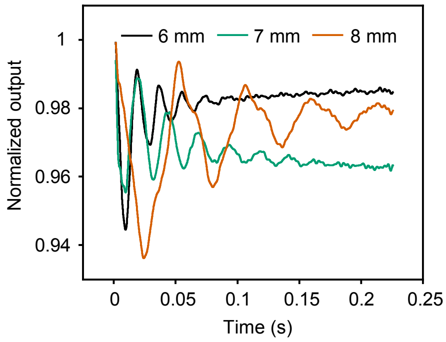

**Fig. S24. Vibration waveforms of the magnetic implants (open cavity) with different diameters in artificial CSF.**

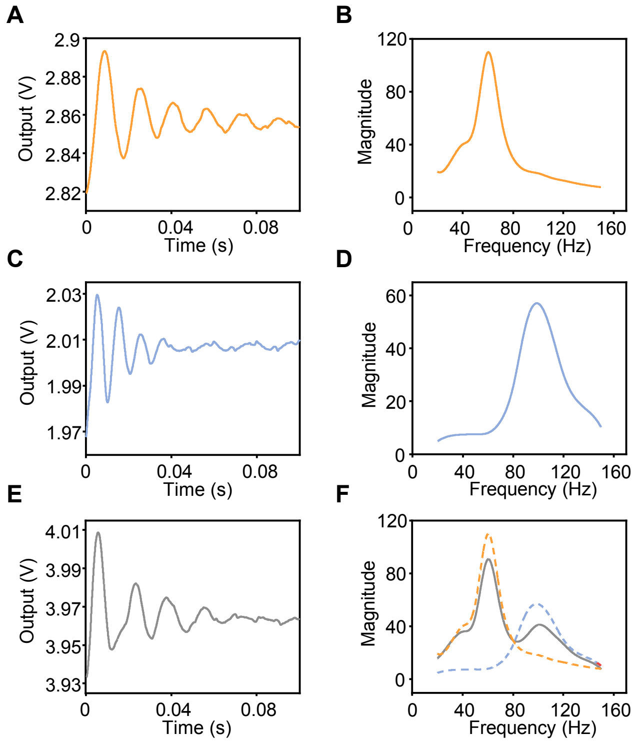

**Fig. S25. Multiplexed sensing using magnetic implants with different vibration frequencies.** (**A** and **B**) Time-domain output voltage (A) and FFT (B) of the vibration waveform of a magnetic implant (open cavity) with a diameter of 8 mm. (**C** and **D**) Time-domain output voltage (C) and FFT (D) of the vibration waveform of a magnetic implant (sealed cavity) with a diameter of 6 mm. (**E** and **F**) Time-domain output voltage (E) and FFT of the signal (F) when two magnetic implants vibrate at the same time.

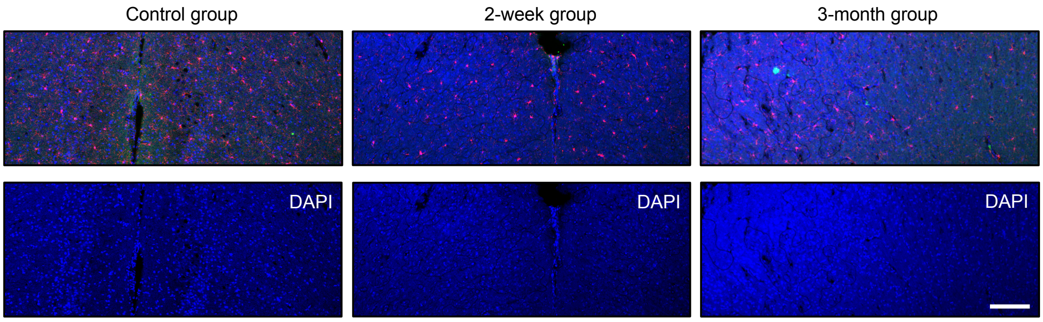

**Fig. S26. Immunofluorescent test.** Immunofluorescent staining images of the control group and cross-sectional areas near the implantation site at 2 weeks and 3 months. Scale bar, 200 µm.

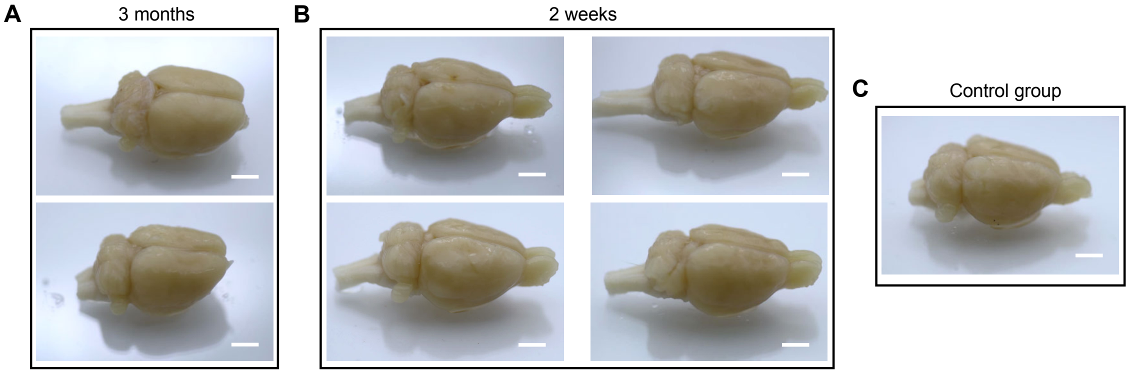

**Fig. S27. Optical images of morphological changes in the rats' brains.** (**A**) 2 weeks after implantation. (**B**) 3 months after implantation. (**C**) Control group. Scale bars, 5 mm.

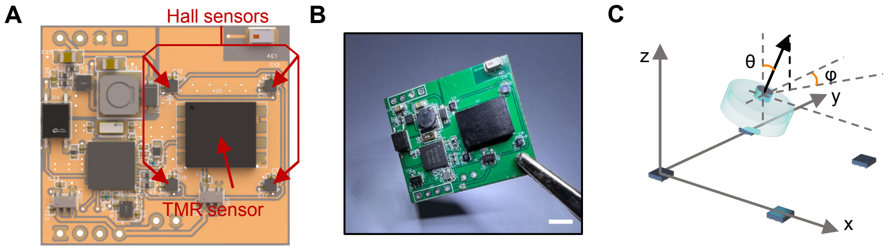

**Fig. S28. Positioning of magnetic implants using 4 Hall sensors.** (**A** and **B**) Schematic illustration (A) and optical image (B) of the circuit in the wearable device. Arrows in red point to the Hall sensors. Scale bar, 5 mm. (**C**) Schematic illustration of the rotation around X axis (θ) and Z axis (φ).

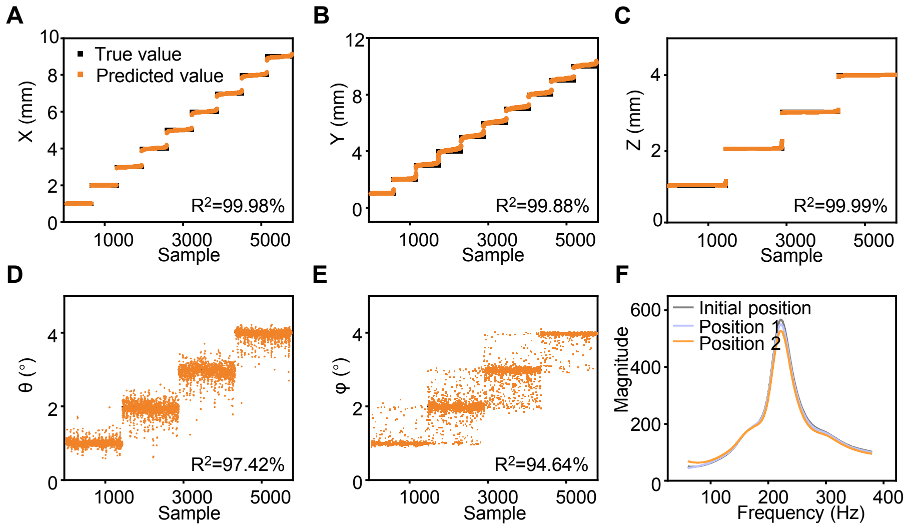

**Fig. S29. Positioning of magnetic implants with MLP regression algorithm.** (**A** to **E**) Relationships between the predicted value and true value for X (A), Y (B), Z (C), θ (D) and φ (E). (**F**) FFT of the vibration of the magnetic implant when actuated by the wearable device at different relative positions. Position 1 has a shift of 0.1 mm along Y axis. Position 2 has a rotation of 0.1° around X axis.

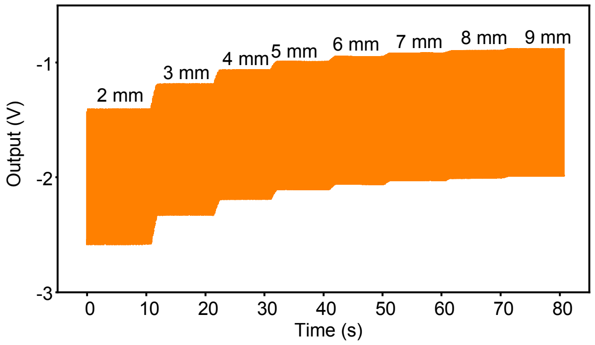

**Fig. S30. Output from the TMR sensor when the wearable device actuates the magnetic implant at different distances.**

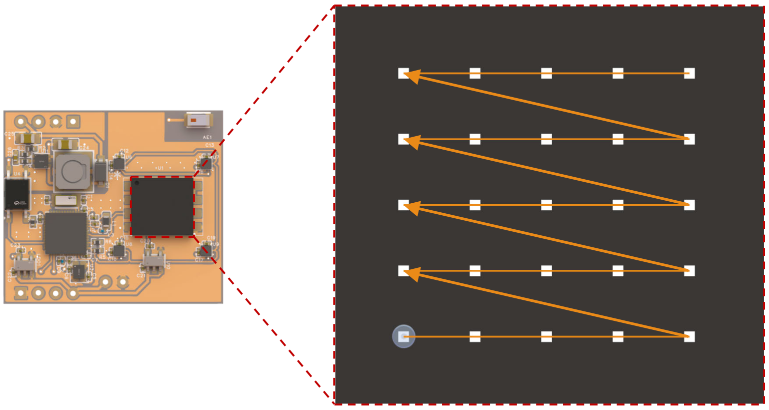

**Fig. S31. Schematic illustration of the moving path for calibration of movement along X and Y axes.**

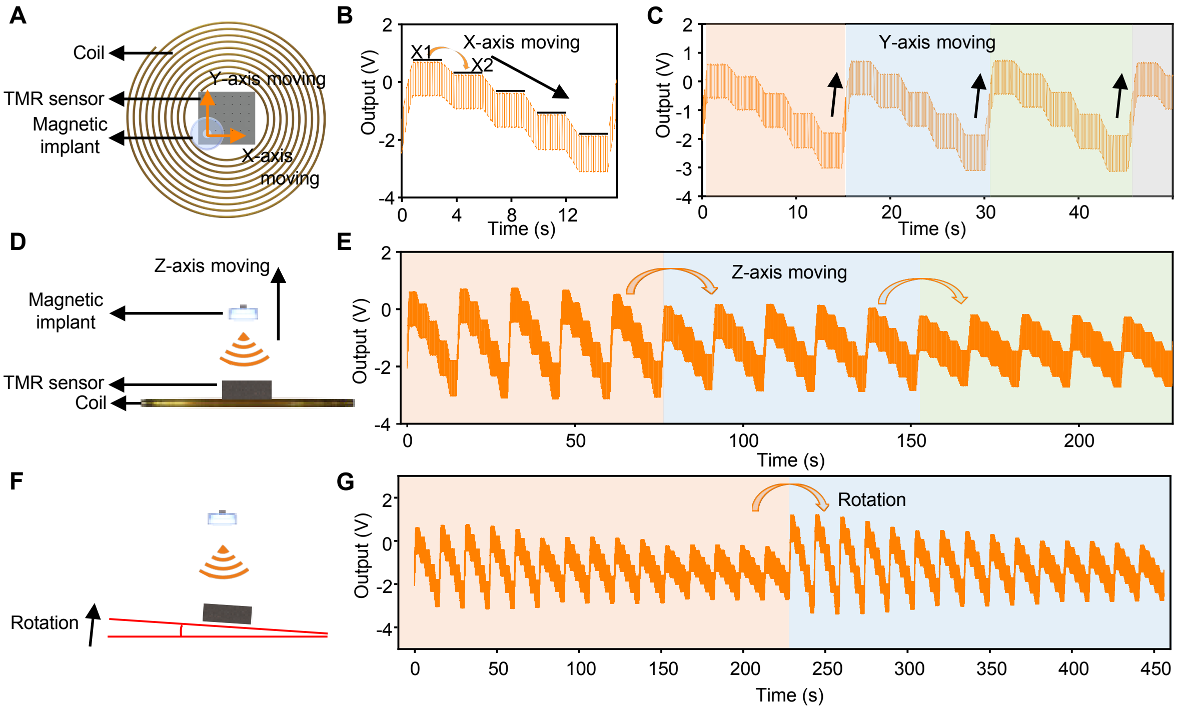

**Fig. S32. Calibration of the wireless pressure sensing at different relative positions.** (**A**) Schematic illustration of the movement along X and Y axes. (**B** and **C**) Output voltage from the TMR sensor when the coil moves along X (B) and Y (C) axes at different positions. (**D**) Schematic illustration of the movement along Z axis. (**E**) Output voltage from the TMR sensor when the coil moves along Z axis at different positions. (**F**) Schematic illustration of the rotation with an angle of θ. (**G**) Output voltage from the TMR sensor when the coil rotates at different angles.

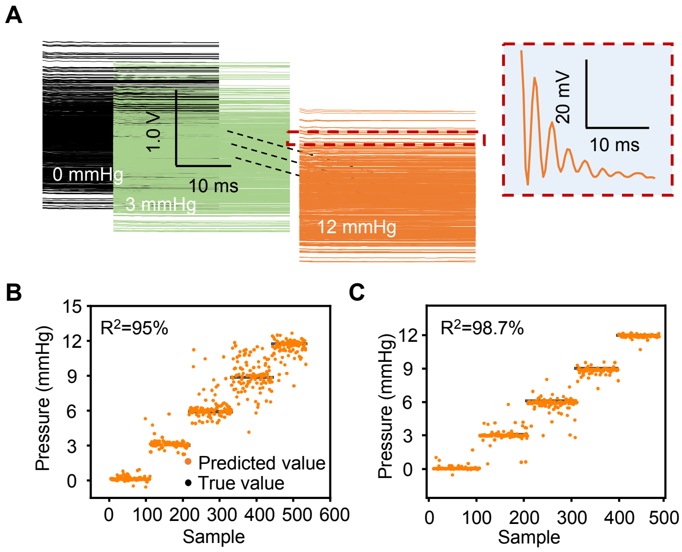

**Fig. S33. Pressure calibration with MLP regression algorithm.** (**A**) Vibration waveforms recorded at all 500 different positions. Waveforms in the same color correspond to the signals recorded at the same pressure condition. The curve inside the red dashed box is the magnified view of one vibration waveform. (**B** and **C**) True pressure (black dots) and predicted pressure (orange dots) at 500 different relative positions, under coarse (B) and fine (C) calibrations. The coarse calibration uses steps of 1 mm for X, Y, and Z, and 1° for θ. The fine calibration uses steps of 0.2 mm for X, Y, and Z, and 0.2° for θ.

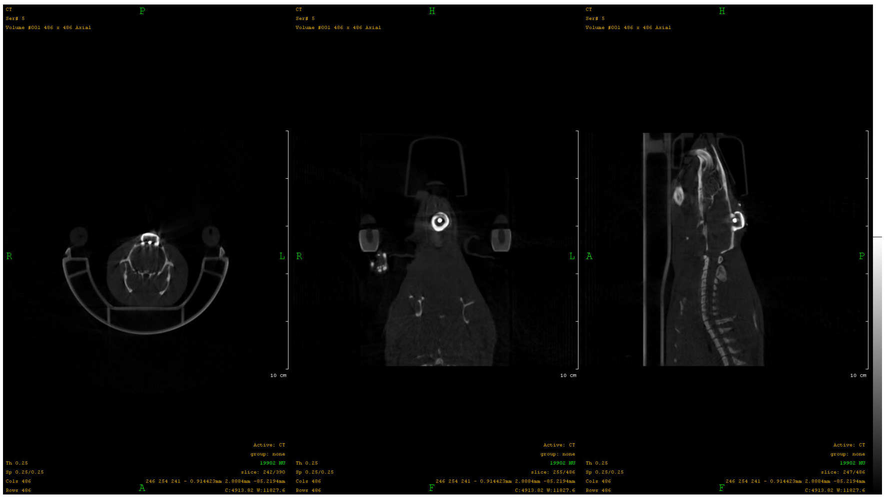

**Fig. S34. CT image of the magnetic implant in rat's brain**.

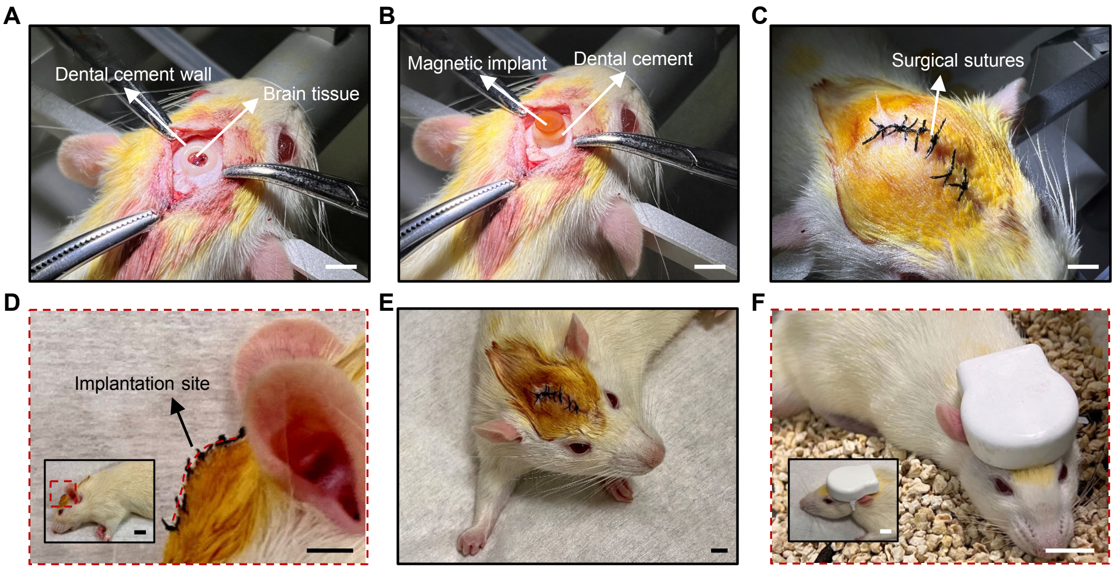

**Fig. S35. Surgical procedures for placing the magnetic implant in rat's brain.** (**A**) A ring of dental cement wall around the craniotomy site. (**B**) Magnetic implant fixed on the wall by dental cement. (**C** and **D**) Different views of the sutured wound at the implantation site. (**E**) An awake rat with the magnetic implant in the brain. Scale bars, 5 mm. (**F**) Optical image of the rat with a mounted wearable device. Scale bar, 1 cm.

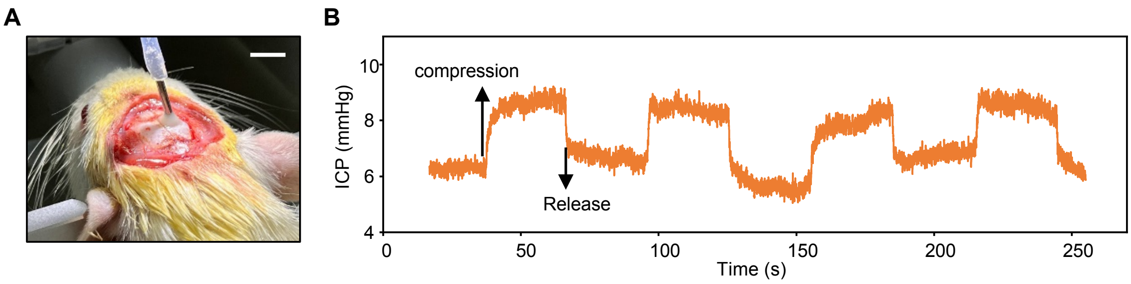

**Fig. S36. ICP measurement from a commercial pressure sensor.** (**A**) Optical image of the commercial pressure sensor implanted on rat's brain. Scale bar, 5 mm. (**B**) Fluctuation of ICP measured from the commercial pressure sensor during abdomen compression.

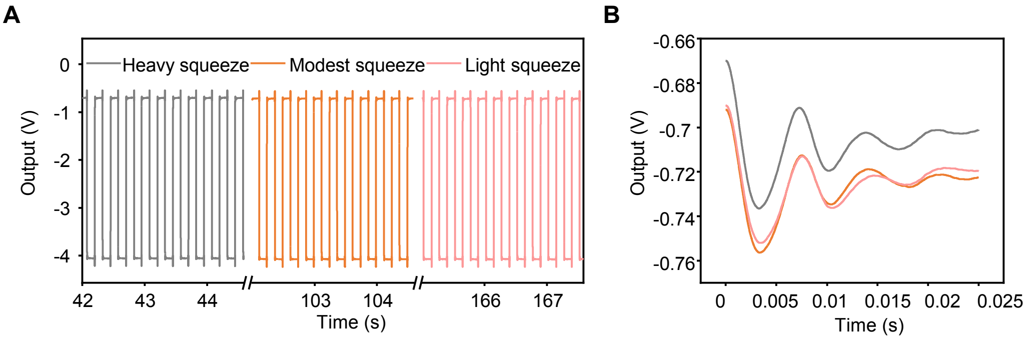

**Fig. S37. Response of the magnetic implant under abdominal compression with various intensities.** (**A**) Waveform captured from the wearable device during the actuation and damped vibration of the magnetic implant under different compression intensities. (**B**) Magnified views of the one-cycle vibration waveforms under different compression intensities.

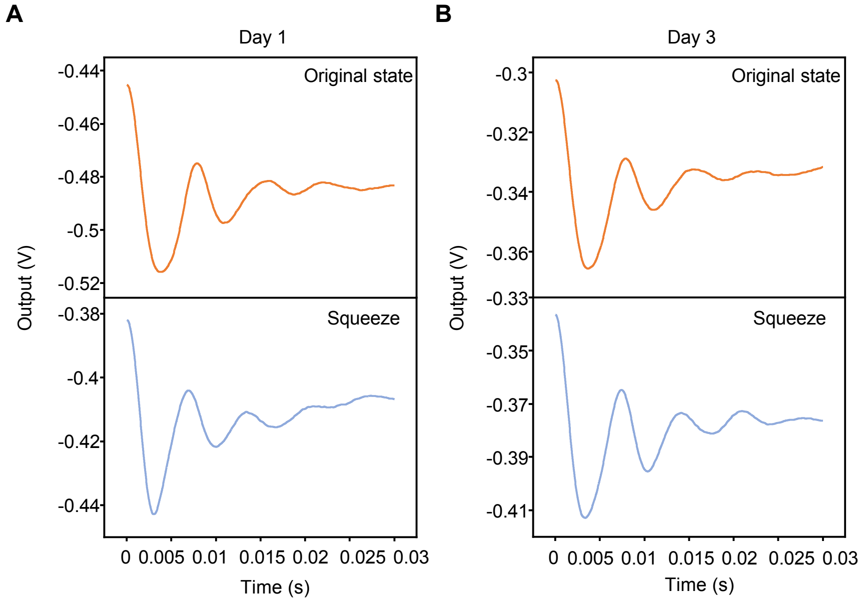

**Fig. S38. Response of the magnetic implant during abdominal compressing on day 1 and day 3.** (**A**) Vibration waveforms with and without squeeze in day 1. (**B**) Vibration waveforms with and without squeeze in day 3.

#### 3. Supplementary Movies:

Movie S1. Two-way wireless communication between the magnetic implants and the fully integrated wearable device.

Movie S2. FEA prediction of the damped vibration of the magnetic implants under different viscosities in liquid environment.

Movie S3. FEA prediction of the damped vibration of the magnetic implants under different pressures in liquid environment.

Movie S4. FEA prediction of the damped vibration of the magnetic implants w/wo absorption in liquid environment.
